# GM-CSF-activated human dendritic cells promote type1 T follicular helper cells (Tfh1) polarization in a CD40-dependent manner

**DOI:** 10.1101/2022.04.28.489850

**Authors:** Sarantis Korniotis, Melissa Saichi, Coline Trichot, Caroline Hoffmann, Elise Amblard, Annick Viguier, Sophie Grondin, Floriane Noel, Hamid Mattoo, Vassili Soumelis

## Abstract

T follicular helper (Tfh) cells are specialized CD4^+^ T cells that regulate humoral immunity by providing B cell help. Tfh1 sub-population was recently identified and associated with severity in infection and autoimmune diseases. The cellular and molecular requirements to induce human Tfh1 differentiation are unknown. Our work investigated the role of human dendritic cells (DC) in promoting Tfh1 differentiation and their physiopathological implication in mycobacterium tuberculosis and mild COVID-19 infection.

Activated human blood CD1c^+^ DC were cocultured with allogeneic naive CD4^+^ T cells. Single-cell RNA sequencing was then used alongside protein validation to define the induced Tfh lineage. DC signature and correlation with Tfh1 cells in infected patients was established through bioinformatic analysis.

Our results show that GM-CSF-activated DC drove the differentiation of Tfh1 cells, displaying typical Tfh molecular features, including 1) high levels of PD-1, CXCR5, and ICOS expression; 2) BCL6 and TBET co-expression; 3) IL-21 and IFN-γ secretion. Mechanistically, GM-CSF triggered the emergence of two distinct DC sub-populations defined by their differential expression of CD40 and ICOS-ligand (ICOS-L), and distinct phenotype, morphology, transcriptomic signature, and function. We showed that Tfh1 differentiation was efficiently and specifically induced by CD40^high^ICOS-L^low^ DC in a CD40-dependent manner. Tfh1 cells were positively associated with a CD40^high^ICOS-L^Low^ DC signature in patients with latent mycobacterium tuberculosis and mild COVID-19 infection.

Our study uncovers a novel CD40-dependent human Tfh1 axis. Immunotherapy modulation of Tfh1 activity might contribute to control diseases where Tfh1 are known to play a key role, such as infections.

**Significance Statement:** Dendritic cells (DC) play a central role in triggering the adaptive immune response due to their T cell priming functions. Among different T cell subsets, it is still not clear how human type1 T follicular helper cells (Tfh1) differentiate. Tfh1 cells are implicated in several physiopathological conditions, including infections. Here we show that GM-CSF induces diversification of human DC. Only CD40^high^ICOS-L^Low^ DC were able to drive Tfh1 cell differentiation. We found that CD40^high^ICOS-L^Low^ DC signature was associated to Tfh1 cells in mycobacterium tuberculosis and COVID-19 patients. Our data reveal a previously undescribed pathway leading to human Tfh1 cell differentiation and highlight the importance of GM-CSF and CD40 as potential targets for the design of anti-infective therapies.

## Introduction

Tfh cells provide critical help to B cells for proliferation, somatic hypermutation, class-switch recombination, and differentiation into antibody (Ab)-producing plasma cells (1,2). They have been associated to several human diseases, including viral and bacterial infections (3-8) asthma (9-11), cancer (12-14) and autoimmune diseases (15-17), but the mechanisms underpinning their development and functions are not well characterized.

Circulating Tfh cells were divided into sub-populations sharing key phenotypic and functional characteristics with other T helper lineages such as Th1, Th2 and Th17 and displaying different capacity in regulating B cell responses (18). The nature of the inflammatory microenvironment affects Tfh differentiation programs, which may subsequently regulate B cell immunity. Tfh1 cells were identified based on their cytokine profile, which is characterized by the co-production of IL-21 and IFN-γ, and specific phenotypic features: PD-1^+^ ICOS^+^ CXCR5^+^ CXCR3^+^ CCCR6^-^(18). Tfh1 cells were increased in HIV infection where the Tfh1 population represents a major fraction of the viral reservoir (19-21). Studies in mycobacterium tuberculosis infection (MTB) have revealed a role of Tfh1 cells and association with disease severity (22-25). Mouse studies showed an important role for Tfh1 in controlling Zika virus infection, as well as Lymphocytic Choriomeningitis Virus (LCMV) infection, mainly through IFN-γ secretion (26,27). Additionally, it has been recently shown that circulating Tfh1 cells (cTfh1) are positively correlated with the magnitude of viral specific antibodies in both influenza and COVID-19 patients (5-7,28,29). In line with the results from the influenza vaccination, recently published findings in mice revealed that immunization with SARS-CoV-2 mRNA elicited potent viral-specific Tfh1 cells needed to produce long-lived plasma cells (29-31).

Tfh differentiation requires cooperation between antigen-specific interactions and signaling pathways, co-stimulation, cytokines, and chemokine receptors (32). Dendritic cells (DC)-derived cytokines, such as IL-6, IL-12, IL-23 and TGF-β promote surface CXCR5 expression on Tfh cells, facilitating their migration towards the T-B cell interface within the secondary lymphoid organ germinal centers (GC) (33,34). Additionally, human Tfh differentiation is driven by Activin-A in a SMAD2 and SMAD3 dependent way (35). Given the established Tfh subset diversity, it is now of critical physio pathological and therapeutic importance to identify cellular and molecular mechanisms controlling specific Tfh differentiation pathways. We have previously shown that TSLP-activated DC could promote Tfh2 differentiation through OX40 ligand (36). However, the factors inducing human Tfh1 differentiation remain elusive.

Here, single-cell RNA sequencing (scRNAseq) analysis of human CD4^+^ T cells differentiated by GM-CSF-activated blood DC (GM-CSF-DC) revealed the presence of bona fide Tfh1 cells. Mechanistically, GM-CSF induced diversification of human DC into two phenotypically, transcriptionally, and functionally distinct subsets. Only CD40^high^ICOS-L^Low^ DC could efficiently drive Tfh1 polarization, in a CD40-dependent manner. Moreover, we found that Tfh1 cells were positively correlated with a signature of CD40^high^ICOS-L^Low^ DC in two different clinical settings of infection, mycobacterium tuberculosis and active COVID-19 patients. Overall, our results define a novel Tfh1 differentiation pathway, with potential molecular targets for its pharmacological manipulation.

## Results

### ScRNAseq reveals a Tfh1 polarization program induced by GM-CSF-activated DC

We decided to revisit human Th cell differentiation induced by GM-CSF-DC in a comprehensive manner using single-cell RNA sequencing (scRNAseq). Primary human naive CD4^+^ T lymphocytes were co-cultured 6 days with allogeneic primary blood cDC2 previously activated for 24h with LPS (LPS-DC), GM-CSF (GM-CSF-DC) or cultured in medium only (Medium-DC). scRNAseq was performed in sorted T cells after six days of co-culture, using the 10X-Genomics platform, on an average 5,600 cells per DC condition. This led to a total of 17,070 high quality sequenced cells, with an average 4,000 detected genes per cell. To probe the dataset for prototypical Th subsets, we used knowledge-driven signatures characteristic of Th1, Th2, Th17, and Tfh cells (Table S1). To be able to dissect Th diversity, we applied those signatures to CD4^+^ T cells generated with GM-CSF-DC. We dimensionality reduced the data using Principal Component Analysis (PCA) and we visualized them using Uniform Manifold Approximation and Projection (UMAP). Th1 and Th2 signatures were enriched in distinct cell clusters, representing 11.6% and 12.0% of cells, respectively (Fig. 1A). The Th17 signature was not enriched in the whole dataset, indicating that GM-CSF-DC do not have the potential to induce Th17 differentiation program. Interestingly, two clusters were enriched in the Tfh signature. One of them co-expressed a Th1 signature, suggestive of Tfh1 cells (11.6%), while the other (16.5%) did not overlap neither with Th1-nor Th2-enriched clusters (Fig. 1A). Tfh-enriched clusters were not detected in T cells activated by either Medium-DC or LPS-DC (fig. S1A). To better define the specific contribution of Tfh markers to the Tfh signature enrichment in the different cell clusters, we represented the expression of four key Tfh genes: BCL6, PDCD1 (PD-1), CXCR5 and IL21. All genes were highly expressed exclusively in CD4^+^ T cells differentiated by GM-CSF-DC (Fig. 1B) (p=0, comparing GM-CSF-DC and LPS-DC, GM-CSF-DC and Medium-DC, permutation test). Interestingly, the Tfh1-enriched cluster displayed higher expression for both the phenotypic markers PD-1 and CXCR5, the transcription factor BCL6 and the key cytokine IL-21, suggesting the induction of a stronger Tfh polarization within Tfh1 cells (Fig. 1C). The induction of Tfh1 cells by GM-CSF-DC was additionally validated by using Pearson pairwise correlation matrices for phenotypic markers, transcription factors (TF) and cytokines known to characterize the three major T helper lineages, Th1, Th2 and Th17. We also observed a high correlation between the cytokines and TF, typical of Th1 and Tfh cells, within the in vitro generated Tfh cells. IL21 expression was strongly associated with Th1-related cytokines IFNG and TNFA, but not with IL4 or IL17A, Th2- and Th17-related cytokines, respectively. Similarly, the master TF of Tfh cells, BCL6, was highly associated with the expression of TBX21, TF of Th1 cells, but not with GATA3, TF of Th2 cells. Additionally, the classical phenotypic markers of Tfh cells, PDCD1 and CXCR5, displayed high correlation with IL-21, IFNG, TNFA, BCL6 and TBX21 but not IL4, IL13, IL17A and GATA3 (Fig. 1D). When the expression level of these genes was represented separately, we observed that GATA3 and IL-4 were exclusively expressed in the Th2-enriched cluster without major expression of IL-13 (fig. S1, B and C). TNFA, IFNG and TBX21 displayed higher levels within the Tfh1-enriched cluster, which highlighted the importance of their co-expression in promoting efficient Tfh1 differentiation (Fig. S1B and C). Overall, our scRNAseq analysis revealed that GM-CSF-DC polarized a significant fraction of naive CD4^+^ T cells into Tfh1 at the transcriptomic level.

**Fig. 1.**
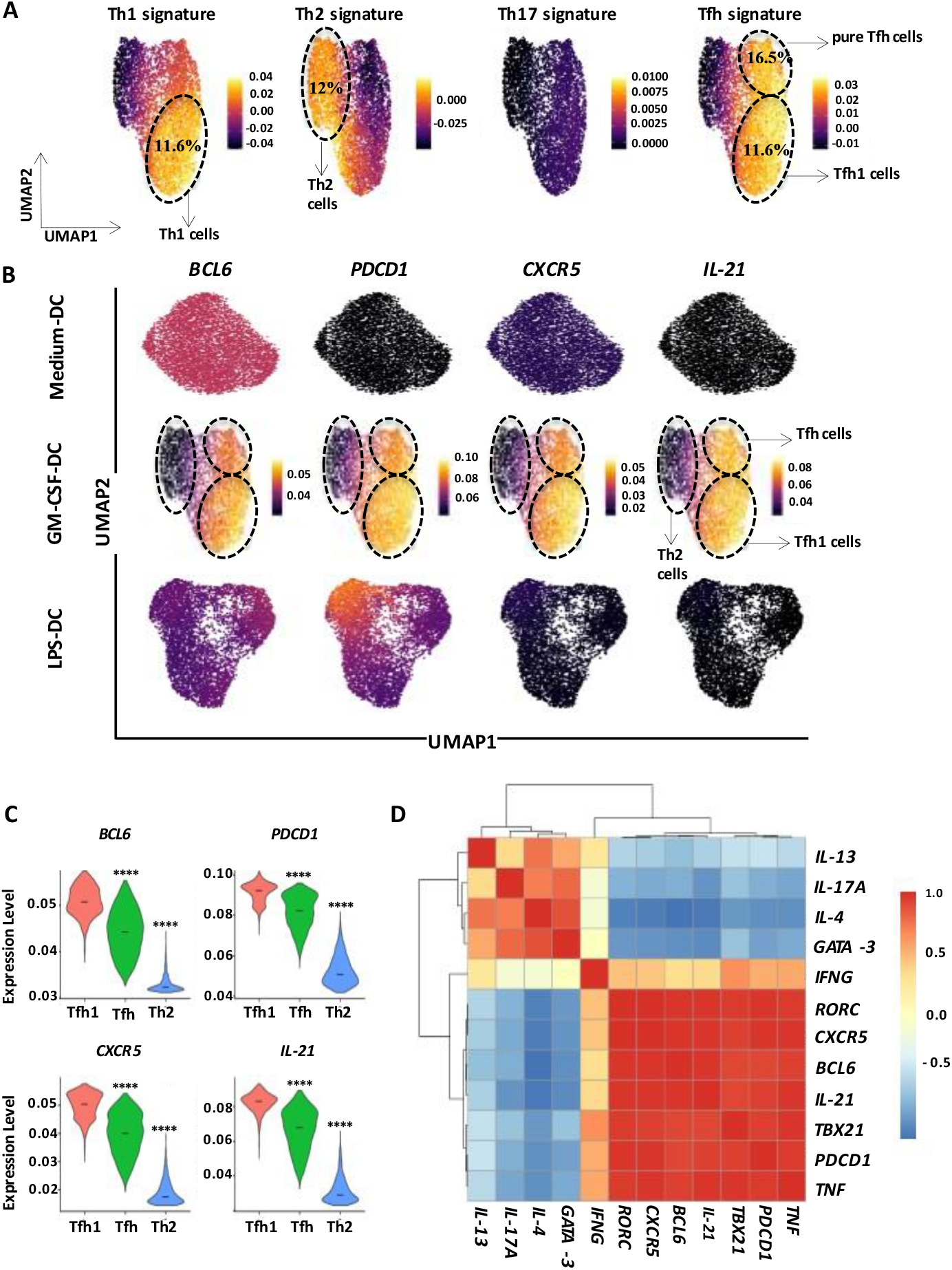
Single-cell RNA sequencing reveals a Tfh1 polarization program induced by GM-CSF-activated DC. **(A)** UMAP plots show the expression scores of Th1, Th2, Th17 and Tfh signatures (Table 1 – Supplementary File) in CD4+ T cells differentiated by GM-CSF-DC. **(B)** UMAP plots of scRNAseq of naive CD4+ T cells differentiated in-vitro by Medium-DC, GM-CSF-DC or LPS-DC. For each plot, the cells were color-coded according to their expression levels of Tfh related genes. **(C)** Violin plot representation of the expression level of BCL6, PD-1, CXCR5 and IL-21 by Tfh-, Tfh1- and Th2-enriched cluster. **(D)** Correlation matrix of the expression values of Tfh, Th1, Th2 and Th17 related genes within CD4+ T cells differentiated by GM-CSF-DC.

### GM-CSF-DC-activated CD4^+^ T cells display a Tfh1 cytokine profile

To validate our findings at the protein level, we characterized the secretion pattern of the GM-CSF-DC-induced T cell population. To do so, we differentially stimulated DC for 24h, and then we cocultured them with allogeneic naive CD4^+^ T cells as for the previous scRNAseq experiment. After six days of co-culture, T cells were re-stimulated with anti-CD3/anti-CD28 (aCD3/aCD28) beads for 24h to measure cytokine secretion in the supernatants, or with PMA/ionomycin and brefeldin-A for 4h, then permeabilized for intracellular cytokine staining (ICS), and analyzed by flow cytometry (Fig. S2A). We found that T cells co-cultured with GM-CSF-DC secreted high levels of CXCL13, IFN-γ, TNF-α and IL-21, as compared to both LPS-DC and Medium-DC (Fig. 2A). The high secretion of IL-21 and CXCL13 confirmed the induction of Tfh-like cells. The concomitant secretion of IFN-γ and TNF-α suggested the possibility of a Tfh1 polarization. GM-CSF-DC-activated T cells produced more IL-9 than LPS-DC-activated T cells, with no significant difference in the secretion of IL-4, IL-10, IL-13, which are prototypical Th2 cytokines, and IL-17A and IL-17F, which characterize the Th17 lineage. (Fig. 2A, and Fig. S2B). These data supported the hypothesis that Tfh1 were being induced by GM-CSF-DC, although at this stage we could not exclude the co-existence of Tfh and Th1 cells within the same T cell population. To investigate this, we studied T cell cytokine co-production at a single cell protein level by ICS. We observed that only GM-CSF-DC induced a high proportion of IL-21-producing T cells (37.2%±1.99) co-producing high levels of IFN-γ (13.46%±1.72) and TNF-α (33.04%±2.89) (Fig. 2B and 2C). We could not identify any cells co-producing IL-21 with either IL-4 or IL-17A, confirming our hypothesis that GM-CSF-DC favored differentiation towards a Tfh1 fate (Fig. 2B, 2C and Fig. S2C). Our results indicated that GM-CSF-DC were also inducers of IFN-γ^+^IL-21^-^ Th1 cells (20.72%±2.27) (Fig. 2B and 2C). Unsupervised t-SNE analysis of IL-21-producing T cells revealed two clusters: the IFN-γ+ population (25.8%), which stands for Tfh1 cells, and the IFN-γ^-^ population (73.9%), which stands for pure Tfh cells (Fig. 2D). This analysis also revealed that both IL-21^+^IFN-γ^+^ and IL-21^+^IFN-γ^-^ were also producing TNF-α to a similar extent. These results validated our scRNAseq finding on the induction of Tfh1 cells by GM-CSF-DC.

**Fig 2.**
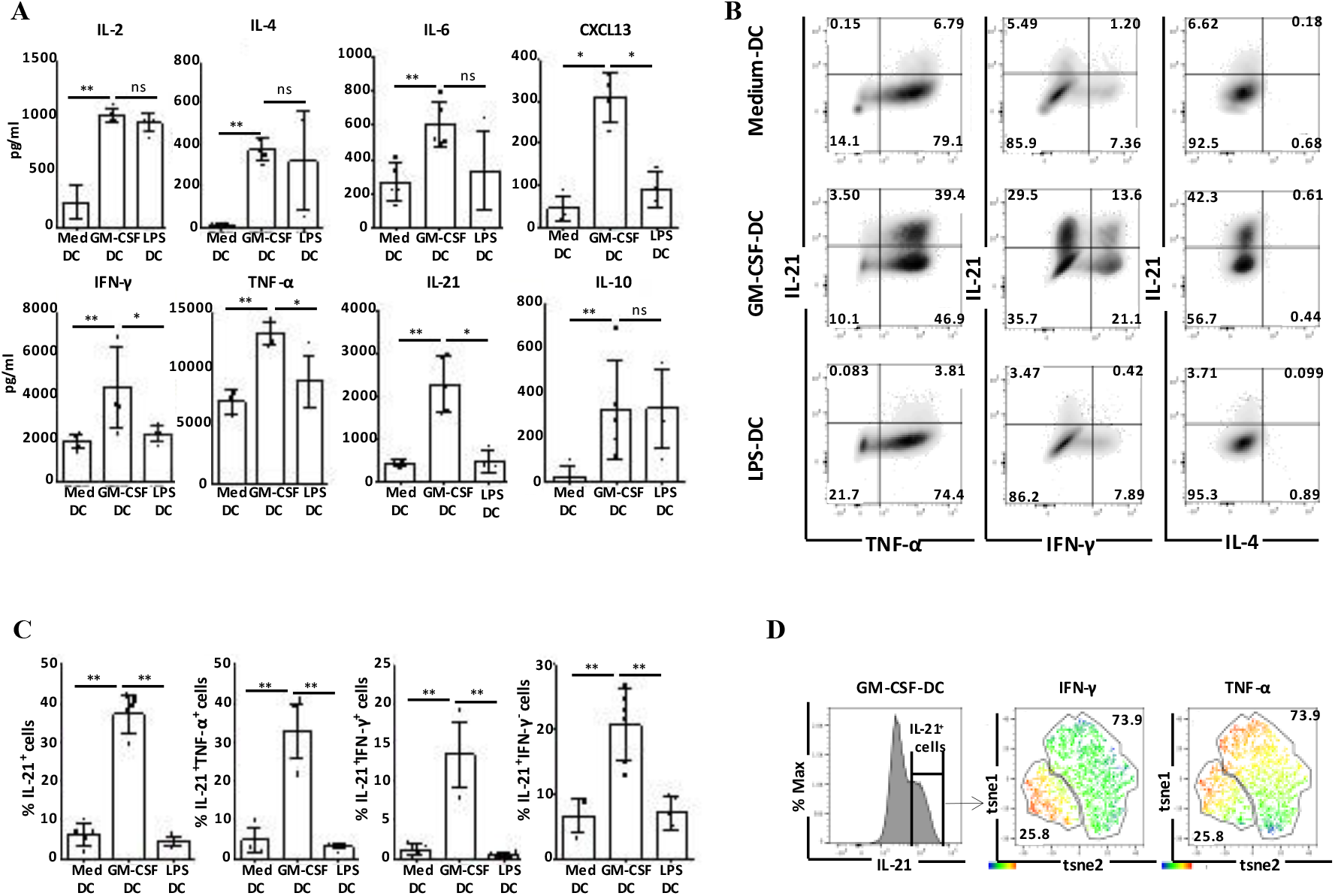
GM-CSF-DC-activated CD4^+^ T cells display a Tfh1 cytokine profile. **(A)** T cell cytokine quantification after coculture of naive CD4^+^ T cells with Medium-DC, GM-CSF-DC or LPS-DC; mean ± SEM from n=5 etc. **(B)** Intracellular FACS staining for T cell cytokines. Plots represent cells gated in CD4^+^ live cells for one representative donor. **(C)** Percentages of IL-21^+^, IL-21^+^TNF-α^+^, IL-21^+^IFN-γ^+^, IL-21^-^IFN-γ^-^ cells as shown in B; mean ± SEM from n=6. **(D)** IL-21 production in CD4^+^ T cells measured by FACS and t-SNE analysis of IL-21^+^ cells identifying two clusters with different expression levels of IFN-γ and TNF-α for one representative donor.

### GM-CSF-DC-induced T cells have phenotypic and functional features of Tfh1 cells

Next, we investigated the phenotype of GM-CSF-DC-activated T cells, at the optimal time-point of day 4 of co-culture (36). Among total activated CD4^+^ T cells in GM-CSF-DC condition, we detected a very significant population of differentiated T cells with a phenotype of Tfh, co-expressing PD-1 with CXCR5 (38.58%±3.84) and PD-1 with ICOS (29.175%±3.08) (Fig. 3A and 3B). Conversely, we observed that both non-activated (Medium-DC) and LPS-DC were much less efficient in inducing the expression of those three Tfh markers (Fig. 3A and 3B). At this time point, we detected three different activated T cell populations based on the expression levels of PD-1 and CXCR5: 1) T^fh-like^ cells, identified as PD-1^high^CXCR5^+^, 2) T^low^ (PD-1^low^) cells characterized by a PD1^low^CXCR5^+^ phenotype, and 3) PD-1^-^CXCR5^-^ double negative population (DN) (Fig. 3C). We examined the expression level of other Tfh markers in these three populations. As expected, BCL6, a TF needed for the development of Tfh cells, was highly expressed exclusively in T^fh-like^ cells. T^fh-like^ cells were also positive for ICOS, SAP and C-MAF, three additional factors used for the identification of functional Tfh cells (40) (Fig. 3C and 3D). We then asked the question whether these T^fh-like^ cells expressed exclusively their master TF BCL6, or additional TF of other Th cell lineages, TBET (Th1), RΟRγt (Th17) and Gata3 (Th2) at the protein level. We observed that a very significant percentage of T^fh-like^ cells co-expressed BCL6 with either TBET (54.08%±5.47) or RΟRγt (55.61%±6.23) (Fig. 3E and 3F). The expression of RΟRγt might be only a result of transient activation of T cells since the expression of this TF was not associated with significant secretion of Th17-related cytokines, as IL-17A and IL-17F, both in supernatants or intracellularly (Fig. S2B and S2C). The percentage of cells co-expressing BCL6 and GATA3 was low (27.40%±2.56), fitting well with the absence of IL-4 secretion. Conversely, in both T^low^ (PD-1^low^) and DN cells, the co-expression of BCL6 with any of these three TF was significantly reduced, confirming that the in vitro induced PD1^high^CXCR5^+^ T^fh-like^ cells were the only population displaying phenotypic features of Tfh cells (Fig. 3E and 3F). Next, we wanted to verify that the production of Tfh1-related cytokines (IFN-γ and IL-21) derived exclusively from the T^fh-like^ population. Sorted T^fh-like^ and T^low^ cells were stimulated with PMA/Ionomycin and Brefeldin A for 4h and stained intracellularly for IL-21, TNF-α, IFN-γ, TBET and BCL6. As expected, only T^fh-like^ cells co-produced IL-21 and TNF-α (Fig. S2D and S2E). Among them, 19.60%±3.80 of cells expressed IFN-γ. Those cells were also positive for both BCL6 and TBET, confirming the Tfh1 polarization. Functionally, we asked whether the GM-CSF-DC-generated T^fh-like^ cells were able to induce the differentiation of memory B cells into plasma cells. Sorted T^fh-like^ and T^low^ (PD-1^low^) cells were co-cultured with both allogeneic naive and memory B cells (Fig. S3A). After 10 days of co-culture, we identified significant proportions of CD19^low^CD38^high^CD27^high^ cells, standing for plasma cells, only in the T^fh-like^ co-culture condition. More specifically, we found that T^fh-like^ cells induced differentiation of memory B cells into plasma cells. The extent of this plasma cell differentiation was comparable to the one obtained with CpG-B-activated memory B cells, clearly showing the ability of T^fh-like^ cells in exerting prototypical Tfh functions. On the other hand, T^low^ (PD-1^low^) cells promoted very low levels of plasma cell differentiation with both naive and memory B cells (Fig. S3A and S3B). These findings confirm that GM-CSF-DC is a new experimental condition allowing the induction of both phenotypical and functional T^fh-like^ cells.

**Fig. 3.**
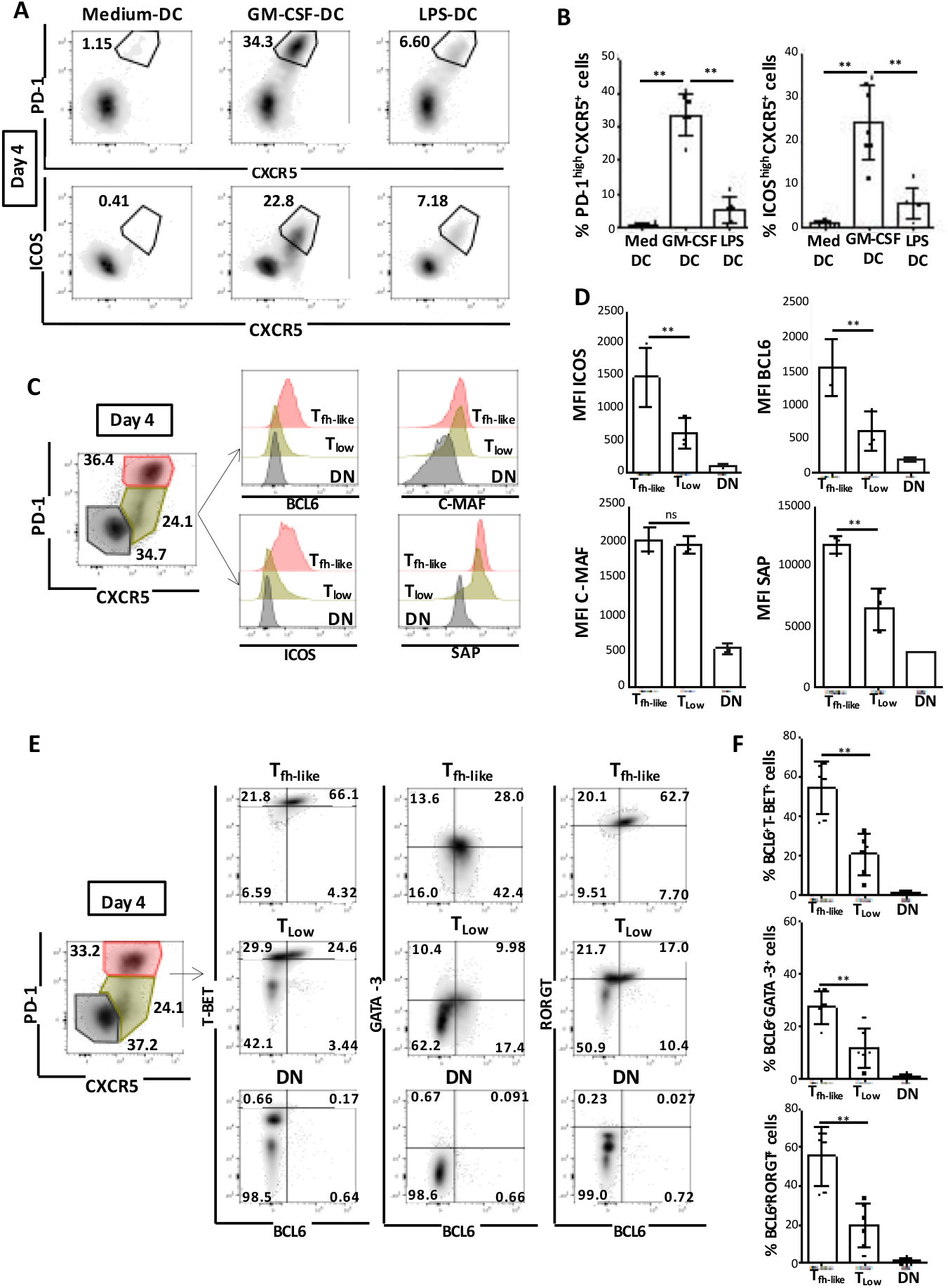
GM-CSF-DC-induced T cells have phenotypic and functional features of Tfh1 cells. (A) PD-1, ICOS and CXCR5 FACS analysis in CD4^+^ T cells after co-culture with Medium-DC, GM-CSF-DC or LPS-DC for one representative donor. **(B)** Percentages of PD-1^high^CXCR5^+^ and ICOSh^high^CXCR5^+^ cells as shown in A; mean ± SEM from n=7. **(C)** Identification of three populations in T cells differentiated by GM-CSF-DC. T^fh-like^ cells (PD-1^high^CXCR5^+^) are shown in red, T^low^ (PD-1^low^CXCR5^+^) in green and Double Negative DN (PD-1^-^CXCR5^-^) in black. FACS histograms show the expression of BCL6, C-MAF, ICOS and SAP in each of the three populations for one representative experiment. **(D)** BCL6, C-MAF, ICOS and SAP MFI quantification as shown in C (n=3). **(E)** Intra-nuclear staining for the expression of BCL6, TBET, GATA3 and RORγT in T^fh-like^, T^low^ and DN cells induced by GM-CSF-DC for one representative donor. **(F)** Percentages of BCL6^+^TBET^+^, BCL6^+^GATA3^+^ and BCL6^+^RORγT^+^ cells from data as shown in E; mean ± SEM from n=6.

### GM-CSF induces two DC populations with different phenotype, morphology, and plasticity

To explore further the mechanism used by GM-CSF-DC to induce Tfh1 differentiation, we sought to characterize the maturation profile of GM-CSF-DC. After 48 hours of activation, GM-CSF induced strong up-regulation of CD40, ICOS-L, CD86 and PD-L1 and intermediate upregulation of CD80, HLA-DR, CD25 and Nectin-II as compared to Medium (Fig. 4A and 4B). The presence of two peaks of expression for some of these markers over time suggested the existence of two DC sub-populations. T-SNE analysis of total GM-CSF-DC at day 1, day 2 and day 3 showed that GM-CSF induced the emergence of two different activated sub-populations from day 2 (Fig. 4C). One population expressed high levels of CD40, and low levels of ICOS-L and PD-L1 and was labeled ICOS-L^Low^; whereas the second population expressed low levels of CD40, and high levels of ICOS-L and PD-L1 and was labeled ICOS-L^High^ (Fig. 4C). Interestingly, the percentage of ICOS-L^Low^ DC was higher than ICOS-L^High^ DC at both day 2 and day 3 of GM-CSF stimulation (45.180%±4.14 at day 2 and 39.45%±4.0 at day 3 for ICOS-L^High^, 54.06%±4.0 at day 2 and 59.12%±2.2 at day 3 for ICOS-L^Low^) (Fig. S4A). For further analysis, we sorted the two sub-populations at day 2 using CD40 and ICOS-L staining (Fig. 4D). Using a Flow-Stream Imaging approach, we detected morphological differences between the two sub-populations. The ICOS-L^High^ DC displayed a typical morphology of activated DC with high FSC/SSC levels, typical dendrites, and very low levels of circularity. The ICOS-L^Low^ DC were rounder, FSC/SSC^Low^, with no dendrites, suggesting a less mature stage of differentiation (Fig. 4E and Fig. S4B). To address the question of cell plasticity, sorted ICOS-L^High^ and ICOS-L^Low^ DC at day 2 were re-stimulated with GM-CSF for additional 24h and 48h. We observed a stable phenotype of ICOS-L^High^ DC after 24h as well as 48h of stimulation (Fig 4F and 4G). On the contrary, ICOS-L^Low^ DC were more plastic since 56.3%±9.9 of cells at 24h and 49.8%±6.0 of cells at 48h acquired a phenotype of ICOS-L^High^ DC (Fig. 4F, 4G, 4H and 4I). We then thought to analyze the expression of GM-CSF receptor a chain (CD116) on the two sub-populations of GM-CSF-DC, as well as in Medium-DC and LPS-DC. ICOS-L^High^ DC displayed higher levels of CD116 compared to ICOS-L^Low^ DC, which could contribute to the lower secondary response of ICOS-L^Low^ DC to GM-CSF exposure (Fig. 4J). We observed that total GM-CSF-DC displayed two peaks of CD116 expression at both day 1 and day 2, suggesting that this marker could be also used to separate the two GM-CSF-induced sub-populations. ICOS-L^High^ (CD116^+^) DC expressed lower levels of GM-CSF-R compared to LPS-DC but comparable levels to Medium-DC, whereas the ICOS-L^Low^ DC (CD116^low^) down-regulated the expression of GM-CSF-R (Fig. S4C). We hypothesized that the emergence of these two sub-populations could also be dose dependent. To address that, we activated DC for 48h with increasing doses of GM-CSF (10ng/ml to 100ng/ml). The emergence of these two sub-populations was dose-independent since the ratio between ICOS-L^High^ and ICOS-L^Low^ DC (10:6) did not vary between the different doses tested (Fig. 4K). These data raised the question whether the GM-CSF-DC-induced sub-populations displayed also functional differences in promoting Tfh polarization.

**Fig. 4.**
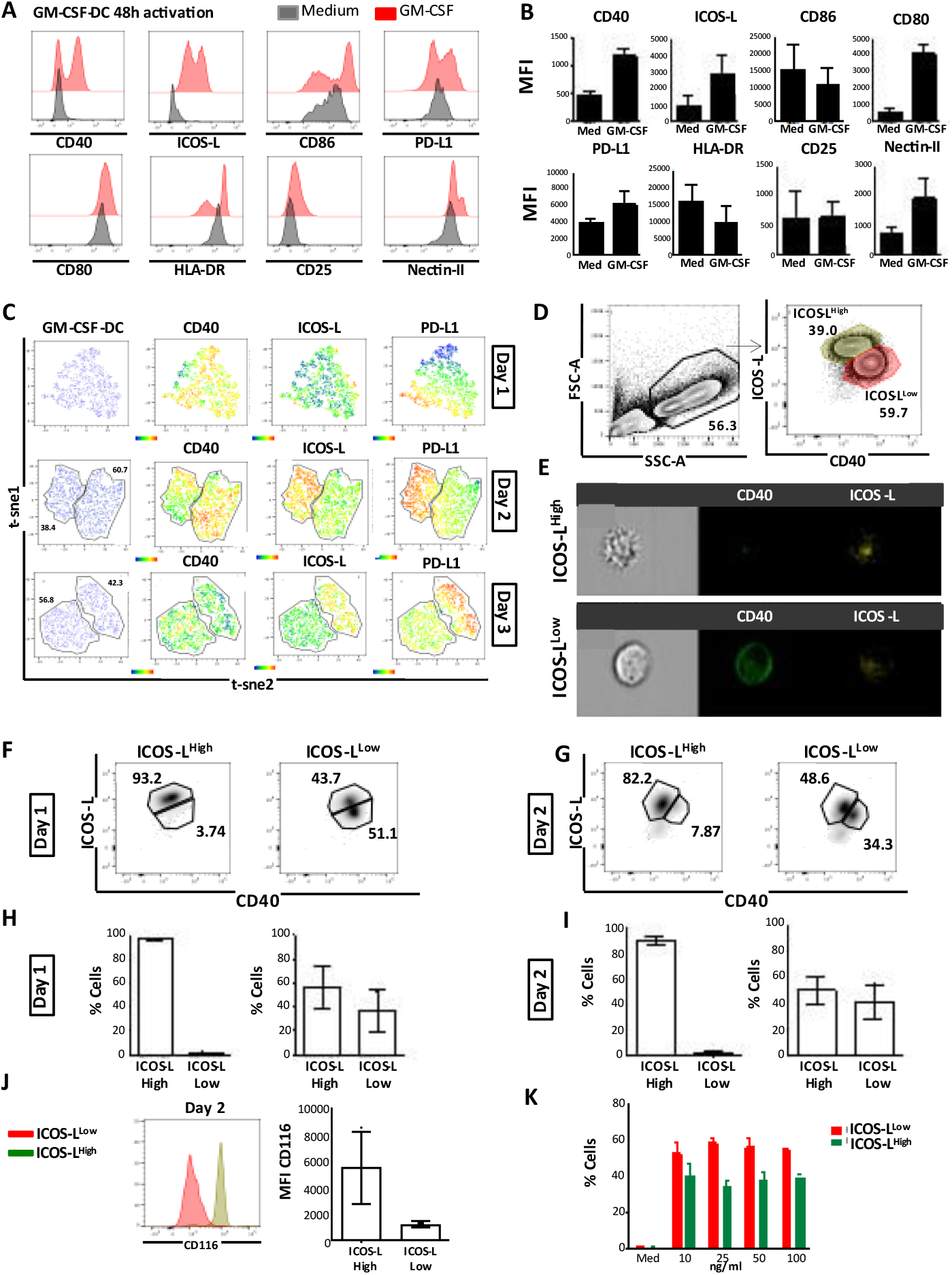
GM-CSF induces the emergence of two populations of DC with different phenotype, morphology and plasticity. **(A)** GM-CSF-DC and Medium-DC phenotypic FACS analysis for one representative donor. **(B)** MFI quantification for data as shown in A (n=6). **(C)** T-SNE analysis of GM-CSF-DC for the identification of different cell clusters based on the expression of CD40, ICOS-L and PD-L1 for one representative experiment. **(D)** FACS analysis for the detection and cell sorting of two different GM-CSF-induced DC sub-populations for one representative experiment. **(E)** Imaging Flow Cytometry analysis of ICOS-L^High^ and ICOS-L^Low^ GM-CSF-DC displaying morphology and expression of CD40 and ICOS-L for one representative donor (n=3). **(F)** and **(G)** ICOS-L and CD40 expression by ICOS-L^High^ and ICOS-L^Low^ GM-CSF-DC after one or two days of GM-CSF re-stimulation for one representative experiment. **(H)** and **(I)** Percentages of ICOS-L^High^ and ICOS-L^Low^ cells as shown in F and G (n=3). **(J)** GM-CSF-Ra chain (CD116) expression in ICOS-L^High^ (green) and ICOS-L^Low^ (red) for one representative experiment; histograms show the mean ± SEM from four independent experiments. **(K)** Percentages of ICOS-L^High^ and ICOS-L^Low^ subsets of DC recovered after 48h of DC activation with different doses of GM-CSF (n=5).

### GM-CSF-induced ICOS-L^Low^ DC promote Tfh1 polarization

We performed 4-day and 6-day co-cultures of sorted ICOS-L^High^ and ICOS-L^Low^ cells with allogeneic naive CD4^+^ T cells. At day 4, we observed that ICOS-L^Low^ DC were more efficient in driving Tfh differentiation since 29.6%±4.6 of cells displayed a Tfh phenotype compared to 13.5%±1.3 of cells in the ICOS-L^High^ DC condition (Fig. 5A and 5B). Within T^fh-like^ cells, ICOS-L^Low^ DC induced higher co-expression of BCL6 with either TBET (75.4%±3.7) or ICOS (42.7%±4.3), as compared to ICOS-L^High^ DC-activated T cells (64.28%±3.69 for BCL6/TBET and 31.8%±2.7 for BCL6/ICOS). GATA3 was also co-expressed with BCL6, but at lower levels (58.4%±3.2 for ICOS-L^Low^ and 61.9%±4.2 for ICOS-L^High^) (Fig. 5A and 5B). The higher percentage of PD1^high^CXCR5^+^ T^fh-like^ cells induced by ICOS-L^Low^ DC together with a significant co-expression of BCL6, ICOS and TBET within this population, suggested that this subset displayed the strongest potential in inducing a Tfh1 phenotypic profile. Next, we studied the cytokine production of T cells activated by each GM-CSF-DC sub-population at day 6, either in supernatants or by ICS. We observed that ICOS-L^Low^ DC-activated T cells secreted high amounts of IL-21, IFN-γ, TNF-α and IL-12p70 whereas ICOS-L^High^ DC-activated T cells secreted more CXCL13, IL-4, IL-10, IL-9 and IL-13 (Fig. 5C and Fig. S5). Additionally, ICS showed that ICOS-L^Low^ DC induced higher co-secretion of IL-21 and TNF-α by T cells (17.5%±3.1) as compared to ICOS-L^High^ DC condition (8.4%±1.5). Among IL-21-producing T cells, ICOS-L^Low^ DC promoted a high production of IFN-γ (46.6%±4.7) but not IL-4 (0.4%±0.1). Conversely, ICOS-L^high^ DC induced significant IL-4-secreting T^fh-like^ cells (2.0%±0.5) but much lower levels of IFN-γ (7.5%±2.5) as compared to ICOS-L^Low^ DC. (Fig. 5D and 5E). Based on the cytokine profile of T cells, we concluded that ICOS-L^Low^ DC were efficient inducers of Tfh polarization, favoring the differentiation into Tfh1, whereas ICOS-L^High^ DC only promoted few Tfh-Th2 cells. There was no significant difference in the T cell polarization induced by total GM-CSF-DC as compared to the one induced by ICOS-L^Low^ DC suggesting that this sub-population dominated over the ICOS-L^High^ DC. This can be explained either by the higher number of ICOS-L^Low^ DC (Fig. S4A) or a cytokine competition, which favored the induction of a Tfh1 profile.

**Fig. 5.**
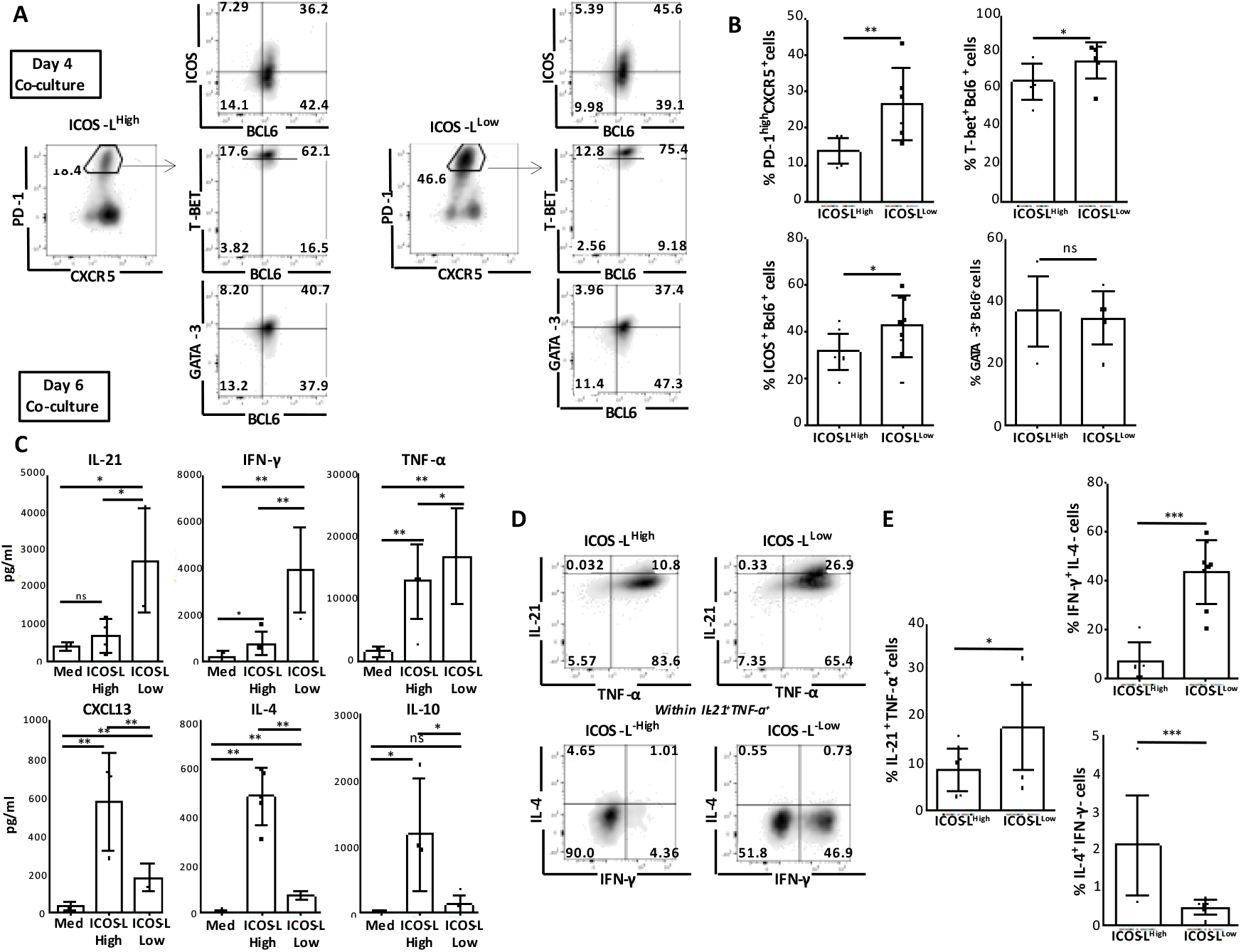
CD40-dependent Tfh1 polarization by GM-CSF-DC. **(A)** BCL6, TBET, GATA3, ICOS FACS analysis in PD-1^high^CXCR5^+^ T cells activated either by ICOS-L^High^ or ICOS-L^Low^ GM-CSF-DC for one representative experiment. **(B)** Percentages of PD-1^high^CXCR5^+^ T cells and ICOS^+^BCL6^+^, TBET^+^BCL6^+^, GATA3^+^BCL6^+^ gated in PD-1^high^CXCR5^+^ T cells from data shown in A; mean ± SEM n=6. **(C)** T cell cytokine quantification after coculture of naive CD4^+^ T cells with ICOS-L^High^ or ICOS-L^Low^ GM-CSF-DC; mean ± SEM n=5. **(D)** Intracellular FACS staining for T cell cytokines. Plots represent cells gated in CD4^+^ live cells for one representative donor. Analysis for IL-4 and IFN-γ is gated in IL-21^+^TNF-α^+^ cells for one representative donor. **(E)** Percentages of IL-21^+^TNF-α^+^ cells and IFN-γ^+^IL-4^-^, IFN-γ^-^IL-4^+^ cells gated in IL-21^+^TNF-α^+^ cells from data as shown in D; mean ± SEM from n=8.

### RNA sequencing reveals major transcriptomic differences between ICOS-L^High^ and ICOS-L^Low^ DC

To further explore the mechanisms by which these two phenotypically, morphologically, and functionally distinct sub-populations of GM-CSF-DC induced Tfh polarization, we performed RNA-sequencing (RNAseq) analysis of sorted GM-CSF-induced ICOS-L^High^ and ICOS-L^Low^ DC (100.000 cells/sample, n=3 donors) (Fig. 6A). We detected 5,118 significantly Differentially Expressed Genes (DEG) between the two sub-populations with an absolute fold-change higher than two. More specifically, 2,414 genes were up regulated in ICOS-L^High^ and 2,704 in ICOS-L^Low^ DC (Fig. 6B and 6C). Focusing on checkpoints and maturation markers, we observed that ICOS-L^Low^ DC expressed more HLA-DRB1 and HLA-DRA, CD276 (B7H3), and CD40, as expected. ICOS-L^High^ DC expressed more negative checkpoints such as CD274 (PD-L1), PDCD1LG2 (PD-L2), TNFRSF9 (4-1BB), IDO1, IDO2, but also the positive checkpoint TNFSF4 (OX40-L), and the maturation molecules CD70, CD80, CD83, CD86 (Fig. 6D). Among secreted molecules, ICOS-L^High^ DC preferentially expressed IL-15, IL-7, IL-32 and the CCR7 ligand CCL19, whereas ICOS-L^Low^ DC expressed more IL-16, IL-1B, TNF, TRAIL, and CCL4 (Fig. 6E). These data raised the question whether some of the DEG were involved in the distinct Tfh polarization programs driven by the two GM-CSF-DC sub-populations.

**Fig. 6.**
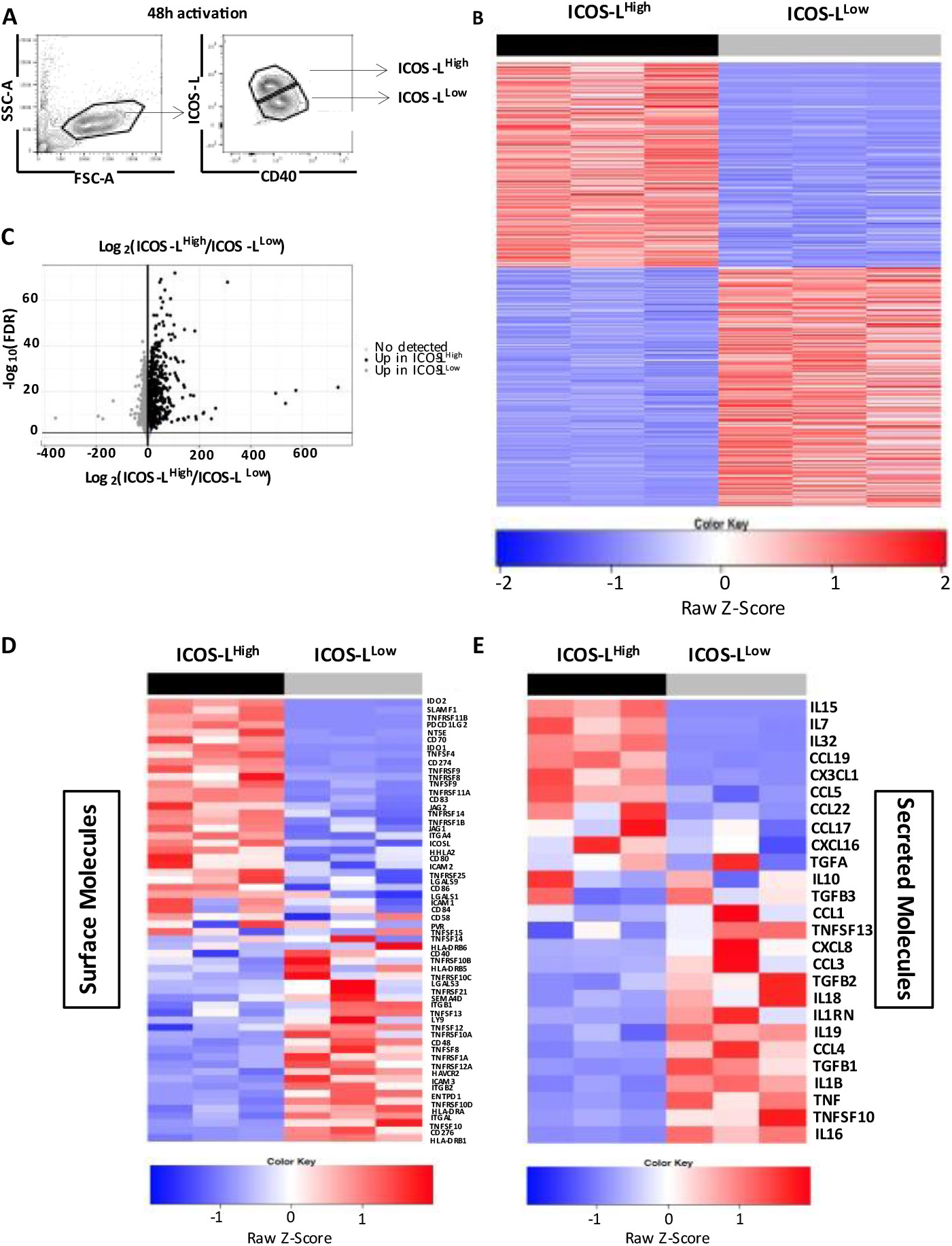
RNA sequencing reveals major transcriptomic differences between GM-CSF-activated ICOS-L^High^ and ICOS-L^Low^ DC. **(A)** Cell sorting strategy for the collection of about 100,000 ICOS-L^High^ and ICOS-L^Low^ DC from 3 independent donors (n=3). **(B)** Heatmap representation of the differentially expressed genes (DEG) between ICOS-L^High^ and ICOS-L^Low^ DC within three donors, z-score values are represented. Up-regulated genes in ICOS-L^High^ are in black; up-regulated genes in ICOS-L^Low^ are in grey, and not differentially expressed genes in light blue. **(C)** Volcano plot of DEG represented through their respective log2 fold-change (X axis) plotted against the FDR (Y axis). **(D)** and **(E)** Heatmap representation of the z-score expression values of DEG from three independent donors for cell surface markers and secreted molecules respectively. Gene expression values were normalized on the B2M and RPL34 housekeeping genes.

### CD40-dependent Tfh1 polarization by GM-CSF-DC

Both CD40 and ICOS-L have been already shown to participate in the crosstalk between DC and T cells (41-43). Since their expression differed in the two sub-populations of GM-CSF-DC, we performed CD40 and ICOS-L blocking experiments to evaluate their respective role in Tfh differentiation. ICOS-L^High^ and ICOS-L^Low^ DC were incubated for 60min with anti-human CD40 and ICOS-L blocking antibodies before the co-culture with allogeneic naive CD4^+^ T cells. CD40 blockade inhibited the induction of a PD-1^high^CXCR5^+^ phenotype in T cells differentiated by both ICOS-L^High^ (No Blocking: 19.8%±2.2, anti-CD40: 9.64%±1.9, p=0.0043) and ICOS-L^Low^ DC (No Blocking: 25.1%±2.9, anti-CD40: 13.5%±2.2, p =0.0098). ICOS-L blockade induced only a slight, but statistically significant reduction of T^fh-like^ cells driven exclusively by ICOS-L^High^ DC (13.0%±2.3 for ICOS-L^High^ and 22.4%±4.4 for ICOS-L^Low^) (Fig. 7A and 7B). We also compared the co-expression of cytokines produced by T cells that were previously polarized by each of the two populations of GM-CSF-DC in the absence of either CD40 or ICOS-L signaling. CD40 blocking reduced the percentage of IL-21^+^TNF-α^+^ cells in both conditions (ICOS-L^High^, No Blocking: 9.3%±2.0, anti-CD40: 4.0%±1.0; ICOS-L^Low^, No Blocking: 17.64%±1.82, anti-CD40: 8.46%±2.90) as well as the percentage of IFN-γ^+^IL4^-^ cells within IL-21^+^TNF-α^+^ T^fh-like^ cells (ICOS-L^High^, No Blocking: 9.3%±1.8, anti-CD40: 4.3%±1.0, p=0.0315; ICOS-L^Low^, No Blocking: 53.21%±5.46, anti-CD40: 33.26%±5.65, p=0.0151). Conversely, CD40 blocking increased the percentage of IL4^+^IFN-γ^-^ cells in both conditions (ICOS-L^High^, No Blocking: 4.6%±0.8, anti-CD40: 14.3%±4.5, p=0.0205; ICOS-L^Low^, No Blocking: 0.41%±0.11, anti-CD40: 3.36%±0.86, p=0.015) (Fig. 7C, 7D, 7E, 7F and 7G). However, ICOS-L blocking did not affect the percentages of IL-21^+^TNF-α^+^ and IL-4^+^IFN-γ^-^ cells in any condition, except for a decrease in IFN-γ^+^IL-4^-^ cells in the ICOS-L^Low^ DC condition (No Blocking: 53.21%±5.46, anti-ICOS-L: 37.30%±4.08, p=0.0409) (Fig. 7C, 7D, 7E, 7F and 7G). These data show a novel and prominent role for CD40 as a new molecule involved in human Tfh1 polarization.

**Fig. 7.**
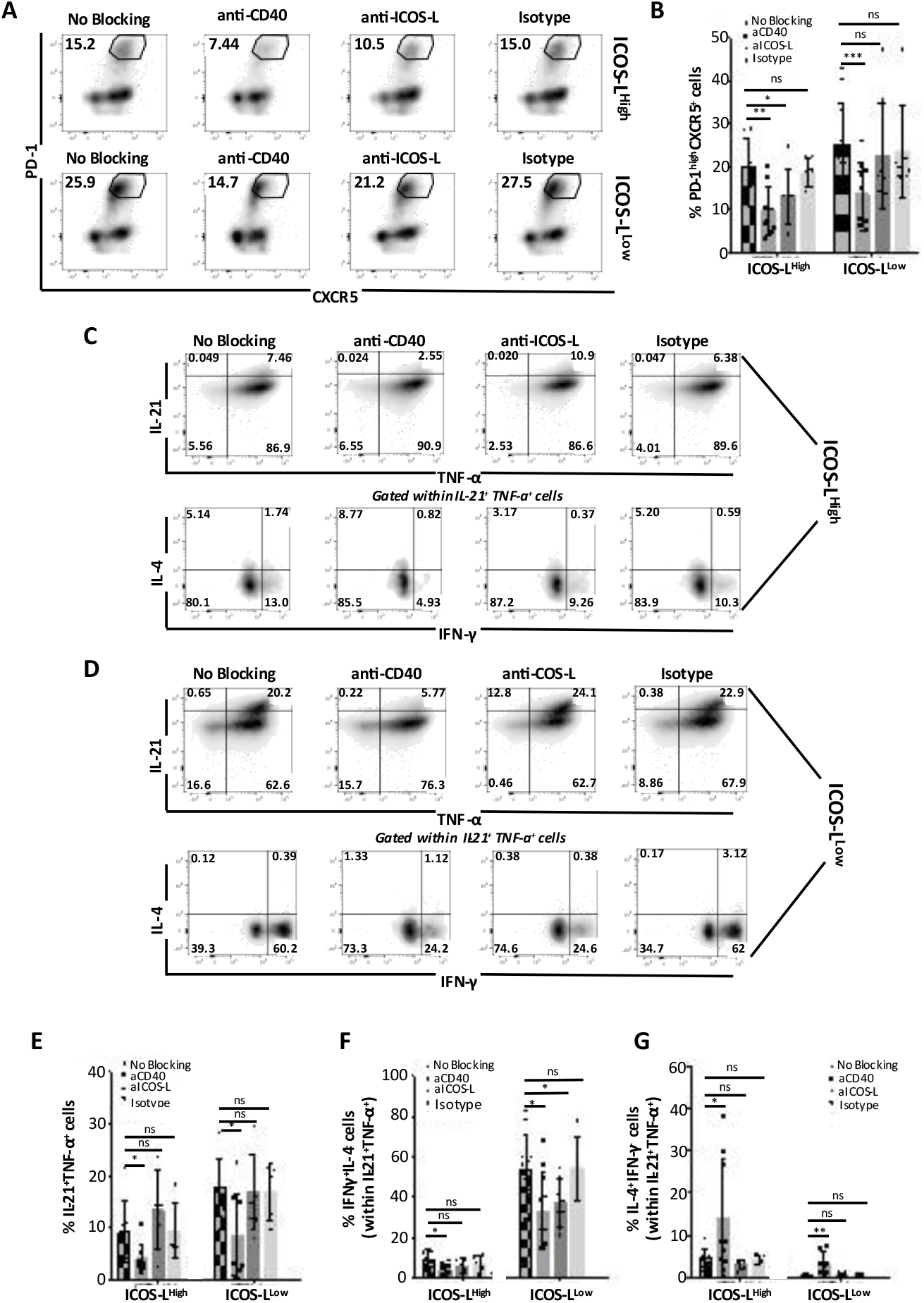
CD40-dependent Tfh1 polarization by GM-CSF-DC. **(A)** PD-1^high^CXCR5^+^ cells gated in total CD4^+^ T cells differentiated by either ICOS-L^High^ or ICOS-L^Low^ DC incubated with blocking antibodies targeting CD40 or ICOS-L and isotype control. One representative experiment is shown. **(B)** Percentages of PD-1^high^CXCR5^+^ cells from data as shown in A; mean ± SEM from n=9. **(C)** and Intracellular FACS staining for T cell cytokines induced either by ICOS-L^high^ or ICOS-L^Low^ DC incubated with blocking antibodies against CD40, ICOS-L and isotype control. One representative experiment is shown. IFN-γ and IL-4 production is gated within IL-21^+^TNF-α^+^ cells. Percentages of IL-21^+^TNF-α^+^ cells, **(F)** IFN-γ^+^IL-4^-^ gated in IL-21^+^TNF-α^+^ cells and **(G)** IL-4^+^IFN-γ^-^ cells gated in IL-21^+^TNF-α^+^ cells from data as shown in D; mean ± SEM from n=8.

### Positive correlation of GM-CSF-induced ICOS-L^Low^ DC with Tfh1 cells in Mycobacterium Tuberculosis and COVID-19

Considering our results, we tested the hypothesis whether GM-CSF-DC sub-populations are associated with the presence of Tfh1 cells in human infections. To explore a possible correlation between GM-CSF-DC signatures and Tfh subsets in vivo, we used two different clinical settings of infection: 1) whole blood microarray data from TB infected patients (38) and 2) scRNAseq of PBMCs from COVID-19 patients (mild, severe, and asymptomatic) alongside severe influenza patients (44). Both studies included healthy controls. Based on our transcriptomic analysis of GM-CSF-DC sub-populations, we selected the top 9-10 genes expressed by each group and we created in-house signatures for ICOS-L^High^ (DC-High) and ICOS-L^Low^ (DC-Low) GM-CSF-DC (Table S2). First, we applied a deconvolution method to assess the presence of all major immune cell types in the microarray data of whole blood from active and latent TB and healthy controls (Fig. S6A), which allowed us to proceed to inter sample and inter cell type comparisons (Fig. S6B). Using Pearson pairwise correlation matrices of DC-High and DC-Low signatures confronted with T effector (Th1, Th2, Th17 and Tfh) cell signatures (Table S3) we observed that Tfh cells were significantly associated only with DC-Low in active and latent TB patient samples, but not in healthy controls (Fig. 8A). Next, we focused on a possible association with the diverse Tfh subsets. Interestingly, we identified a positive correlation between DC-Low and Tfh1 signatures in latent (r = 0.44), but not in active TB, and a negative correlation between DC-High and Tfh1 in both latent (r = 0.54) and active TB (r = 0.50) (Fig. 8B). Neither DC-High nor DC-Low signatures had any significant positive correlation with other subsets of Tfh cells (Tfh2 or Tfh17) (Fig. S6C and S6D) emphasizing the important relation between the sub-population of ICOS-L^Low^ GM-CSF-DC and Tfh1 cells, which might be needed to provide a protective environment in latent TB.

**Fig. 8.**
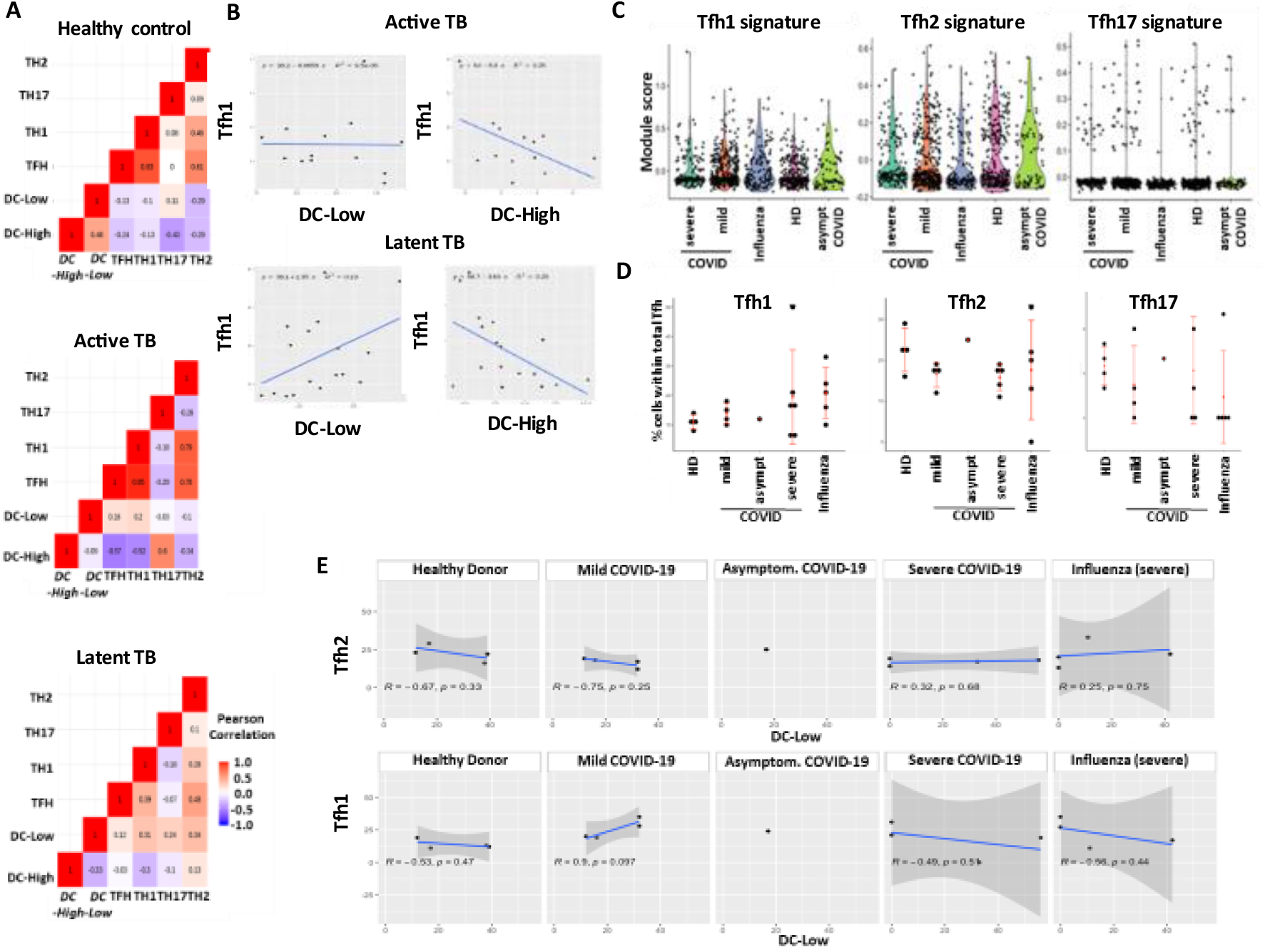
GM-CSF-induced ICOS-L^Low^ DC positively correlate with Tfh1 in Mycobacterium Tuberculosis and COVID-19 infections. **(A)** Pearson pairwise correlation matrices of DC-high and DC-low signatures, confronted with T effector cell signatures for each TB patient group. **(B)** Collinearity assessment of the Tfh1 signature with the DC-High and the DC-Low signatures respectively within Active and Latent TB patients. Each dot represents a sample. A linear regression model was applied to fit the dots repartition. **(C)** Violin plot for the expression level of Tfh1-, Tfh2- and Tfh17-related signatures expressed by all COVID-19 patient groups. **(D)** Percentages of Tfh1, Tfh2 and Tfh17 cells within total Tfh cells in each COVID-19 patient group. No statistical significance was observed for any of these conditions. **(E)** Collinearity assessment of the corresponding signatures of Tfh2 and Tfh1 with the DC subtype signatures DC-Low in all COVID-19 patient group.

In addition, we tried to validate our new experimental findings in a clinical setting of COVID-19 infection (Fig. S7A). We sought to figure out whether we could detect any positive correlation between Tfh1 cells and DC-Low. First, the use of CD3 and CD4 as universal markers for the identification of T cells allowed us to detect them in all the disease groups. We also recovered the T-helper cell subsets using in-house constructed signatures (Fig. 7B and Table S3). Th1, Th2 and Tfh signatures were observed in sufficient numbers in all patient groups whereas very few cells expressed Th17 signature. Interestingly, mild COVID-19 patients revealed higher Th1 numbers as compared to both severe COVID-19 and influenza. The same trend was observed for both Th2 and Tfh cells, suggesting a more efficient adaptive immune response in mild COVID-19 patients (fig. S7C). Since the focus of our study was on Tfh subsets, we used again in-house constructed signatures to identify Tfh1, Tfh2 and Tfh17 cells within the Tfh cluster. The distribution of the corresponding module scores revealed higher positivity for Tfh1 as compared to both Tfh2 and Tfh17 cells (Fig. 8C). This observation was additionally confirmed by statistical comparison of the percentage of positive cells for each signature among total Tfh cells at the patient level. Tfh1 cells are increased mostly in mild COVID-19 patients (Fig. 8D). However, the small cohort size did not allow the conduction of statistically significant tests. In parallel, we looked for the two DC signatures in CD11c^+^BDCA1^+^ cells (Fig. S7D). We could not detect any cells expressing the DC-High signature in all disease groups whereas cells displaying DC-Low signature were present (Fig. S7D). The violin plot representation revealed that enough DC-Low positive cells could be mostly detected in mild COVID-19 patients even if the ratio within the total DC cells is almost the same in all disease groups (Fig. S7E). The estimation of both DC and Tfh subsets is defined by the ratio detected in each patient. Finally, we sought to correlate the percentages of Tfh subsets and DC-Low cells in each disease severity group by applying Pearson correlation test associated to p-value. The only condition where we detected a statistically significant (p<0.1) positive correlation is in mild COVID-19 patients and only between Tfh1 and DC-Low signatures. (Fig. 8E). This observation might be explained by the better response of these patients which need a more activated Tfh1 profile for a more efficient differentiation of plasma cells. The presence of GM-CSF-ICOS-L^Low^ DC might be necessary for inducing efficient Tfh1 responses.

## Discussion

In this study, we provided evidence for the key role of GM-CSF-DC and CD40 in inducing polarization of human naive CD4^+^ T cells into bona fide Tfh1.

BCL6 was shown to be the lineage-defining transcription factor of Tfh cells regulating their functional properties (45-48). Co-expression of BCL6 with other Th transcription factors may imprint Tfh cells with additional functions playing a crucial role in regulating B-cell induced immunity. More specifically, it was shown that viral infections promote TBET expression in Tfh cells, which contributes to IFN-γ secretion, and the type of antibodies produced by plasma cells (26,49). Even a transient TBET expression is sufficient to make Tfh cells secrete IFN-γ for long time (50). We had previously shown that TSLP-DC induce polarization of human naive CD4^+^ T cells into Tfh cells expressing high levels of GATA3 and secreting significant amounts of IL-4 (36). More recently, a new study provided evidence for the existence of Tfh cells (TFH13) with an unusual cytokine profile (IL-13^high^IL-4^high^IL-5^high^IL-21^low^) co-expressing BCL6 and GATA3, affecting high affinity IgE production, and subsequent allergen-induced anaphylaxis (10). In support of these findings, we demonstrated that GM-CSF-DC-induced Tfh cells secreted large amounts of IFN-γ, and expressed high levels of TBET, confirming a strong relationship between transcription factors and subsequent cytokine secretion. However, the identification of two distinct sub-populations of GM-CSF-DC with different functions on T cell polarization raised the question whether the cytokine production can be dissociated from transcription factors. ICOS-L^High^-induced Tfh cells secreted higher levels of IL-4 and much lower amounts of IFN-γ as compared to ICOS-L^Low^-induced Tfh cells, but without any major differences in the expression of either GATA3 or TBET. These observations suggest that the expression of transcription factors might not be always sufficient to identify distinct Tfh subsets since the cytokine profile could reflect a different polarization program. This is not very surprising since ICOS-L:ICOS interactions between the Tfh and germinal center B cells or Tfh and DCs can lead to cytokine secretion that can signal through the Tfh leading to a transcription factor aside from GATA3 and/or TBET. We cannot exclude the possibility that our in vitro system might lead to a transient activation of other transcription factors. Further in vitro and ex vivo studies are needed to better understand the role of transcription factors in the functions of distinct Tfh subsets.

GM-CSF was among the first cytokines shown to efficiently promote DC development in vitro from monocytes and hematopoietic progenitor cells (51-54). It is considered as a regulator of granulocyte, monocyte and DC lineage at all stages of maturation, with effects on cytokine secretion, cytotoxicity and antigen presentation capability (55-57). In our study, we used single cell approaches to show that GM-CSF induces the diversification of human blood DC into distinct sub-populations with different phenotype, morphology, transcriptomic signature, and function. Interestingly, the function of total GM-CSF-DC in activating CD4^+^ T cells was like the effect of ICOS-L^Low^ DC sub-population, showing that the latter had a dominant role in T cell polarization. Previous studies of bulk GM-CSF-DC may have been biased by the function of dominant DC sub-populations and hindered an underlying functional heterogeneity.

CD40 expression on DC was shown to be necessary for promoting survival, cytokine production, and activation of naive T cells (58,59). CD40 is known to induce secretion of IL-12 resulting in enhanced Th1 immune responses (60-62), but there is no direct link with other Th differentiation programs. Interestingly, the reduced Tfh cells in patients with immune deficiencies caused by mutations in CD40 ligand (63) raised the question about a potential role of CD40 in both the generation and maintenance of Tfh cells. In our study we showed by both transcriptomic and single cell protein approaches that CD40 was highly expressed by ICOS-L^Low^ DC, the sub-population of GM-CSF-DC that induced a strong Tfh1 responses. Blocking of CD40 in GM-CSF-DC resulted in reduced frequencies of the common Tfh phenotypic markers PD-1 and CXCR5, together with decreased levels of their key effector cytokine, IL-21. Additionally, inhibition of CD40 expression induced a dramatic decrease in IFN-γ by IL-21-producing Tfh cells, favoring the secretion of IL-4. Altogether, these data suggest that the expression of CD40 by GM-CSF-DC is involved in both Tfh and Tfh1 polarization programs. Its absence might allow other co-stimulatory molecules to dominate their function creating a microenvironment with different effects on the type of Tfh differentiation. CD40 could be considered as a new therapeutic target for the manipulation of Tfh/Tfh1 cells.

The identification of MTB-specific IL-21^+^IFN-γ^+^ Tfh1-like cells (23) together with decreased frequencies of Tfh cells detected in the blood of active patients (24,25) raised the question whether Tfh1 cells display immune regulatory function. Additionally, the protective role of GM-CSF has been already shown in several MTB studies (64-66). We observed a strong positive correlation between DC displaying the ICOS-L^Low^ GM-CSF-DC signature and Tfh1 cells, exclusively in patients with latent form of MTB infection. These results first validate our in vitro findings about the dependence of Tfh1 differentiation on a specific subset of GM-CSF-DC and secondly, open new perspectives for the mechanism used by GM-CSF to induce protective effects in MTB infection. Recent studies demonstrate a strong positive correlation between circulating Tfh1 cells and the magnitude of viral specific antibodies in COVID-19 convalescent patients, highlighting a new crucial role for Tfh1 cells in the fight against viral infections (5,6,67). The severity of COVID-19 infection might be also associated with the function of Tfh1 cells since it has been recently shown that in active severe COVID-19 patients there is a loss of BCL6^+^ Tfh cells and GC, together with an increase in TBET^+^ Th1 cells (68). The huge production of cytokines in severe COVID-19 patients might block GC development, inhibiting the transformation of Th1 cells into Tfh cells (68,69). Taking advantage of publicly available data (70) we identified a positive correlation between Tfh1 cells and ICOS-L^Low^ GM-CSF-DC exclusively in mild COVID-19 patients. Those patients are characterized by a better response to SARS-COV-2; by avoiding a strong cytokine storm, their immune system might allow to an efficient generation of Tfh cells and GC development. Additionally, their Tfh responses could be enhanced by the presence of DC displaying a specific activation profile favoring the polarization of Tfh1 cells. SARS-CoV-2 can make host cells secrete GM-CSF, which could enhance the ability of DC to better prime naive T cells during antigen-specific immune responses (71,72). Additionally, the capacity of GM-CSF to maintain pulmonary function and lung cell-mediated immunity, together with its protective functions in mouse models of influenza, suggested that GM-CSF administration is a possible therapy against COVID-19 (73-75). Indeed, several clinical trials are already ongoing (73).

## Materials and Methods

### DC activation

DC were cultured in RPMI 1640 Medium Gluta-MAX (Life Technologies) containing 10% Fetal Calf Serum (Hyclone), 100 U/ml Penicillin/Streptomycin (Gibco), MEM Non-Essential Amino Acids (Gibco), and 1 mM NA pyruvate (GIB CO). DC were cultured at 106 cells/ml in flat bottom plates for 24h or 48h in the presence of 50 ng/ml rhGM-CSF (Prospec) (unless differently specified) or 100 ng/ml ultrapure LPS (InvivoGen).

### DC/T co-culture

For co-culture, activated-DC were washed twice in PBS 1X and put in culture with allogeneic naive CD4^+^ T cells (10^4^ DC and 5×10^4^ T) in X-VIVO 15 media (LONZA) for the indicated time. For co-culture, CD4^+^ T cells were freshly purified from PBMC the day after DC purification or 2 days later, depending on the experimental condition. Coupling exclusively a single DC donor with a single CD4^+^ T cell donor was used to perform each co-culture experiment. DC were stimulated either with rhGM-CSF or LPS for 24h to activate total cells or only with rhGM-CSF for 48h/72h to induce the emergence of two different sub-populations. After 2 days of activation, sub-populations of DC were electronically sorted (ARIA III, BD) based on the expression of ICOS-L and CD40. Total activated DC or both sub-types of GM-CSF-DC were put in co-culture with allogeneic naive CD4^+^ T cells with the same ratio as mentioned before.

### Blocking Experiments

DC were incubated at 37°C with 50 ng/ml anti–human CD40 antibody, or 50ng/ml anti-human ICOS-L or 50ng/ml of the corresponding isotype control (Biolegend). After 60 min, CD4^+^ naive T cells were added to the culture. Antibodies were maintained for the duration of the co-culture. At indicated time points, cells were either FACS sorted or used for surface or intracellular staining or washed and reseeded at 10^6^ cells/ml and re-stimulated with aCD3/aCD28 beads (LifeTech) for 24h, after which supernatants and cells were collected for analysis. At day 4, T cells were counted and analyzed for the induction of a Tfh profile. At day 6, T cells were counted and analyzed for their cytokine production by FACS.

### Transcriptomic Analysis

Samples were sequenced at QuickBiology, Pasadena CA. Briefly, RNA integrity was checked by Agilent Bioanalyzer 2100. Library for RNA-Seq was prepared according to KAPA Stranded mRNA-Seq poly(A) selected kit with 201-300bp insert size (KAPA Biosystems, Wilmington, MA) using 250 ng total RNAs as input. Library quality and quantity were analyzed by Agilent Bioanalyzer 2100 and Life Technologies Qubit 3.0 Fluorometer. 150 bp paired-end reads were sequenced on Illumina HiSeq 4000 (Illumnia Inc., San Diego, CA). The reads were first mapped to the hg19 UCSC transcript set using Bowtie2 version 2.1.0 and the gene expression level was estimated using RSEM v1.2.15. Downstream analyses were performed using R (v3.6.0) and DESeq2 package (v1.26.0). Differentially expressed genes were determined with an absolute log-fold change threshold at 2 and an adjusted p-value below 0.01

### Microarray Data analysis

We loaded the normalized dataset using GEOquery R package, and GSE19904 as studyID. We selected the top 9 differentially expressed genes between ICOS-L^High^ DC and ICOS-L^Low^ DC from the previous bulk transcriptomic analysis and used the T subtypes signatures previously used. We estimated the fraction of detected cell types in the samples using Quanti Seq deconvolution algorithm from the immunedeconv R package. For each DC and CD4^+^ T cells subtype, we calculated a signature score as the median expression values of the set of genes of a given signature. To assess the relationships between DC subsets and Tfh-Th1 polarization, we plotted the corresponding subtype signature scores and fitter the scatter graphs using linear regression (lm() function on R).

### scRNAseq data analysis of COVID-19 and severe Influenza patients

We performed the analysis on R (version 4.1) using the Seurat R toolkit package for single cell data analysis. Associated metadata of the whole analyzed cells were loaded and added to the Metadata slot of the Seurat object. First, we created a subset Seurat object containing only CD4^+^ T cells. Next, we defined T helper (Th) subsets (Th1, Th2, Th17 and Tfh) using as input our in-house signature genes, and constructed scores for each Th subset for each individual cell, using “AddModuleScore” Seurat function, setting both the number of bins and control genes to n=100. A similar procedure was applied to retrieve CD1c^+^ DC, using the expression levels of canonical markers (positive values for CD1C and ANPEP genes, and absence of expression of THBD gene). Similarly, we applied both DC-Low (ICOSL^Low^ GM-CSF-DC) and DC-High (ICOS-L^High^ GM-CSF-DC) signatures on the DC and followed the same procedure to estimate the percentage of DC for each patient. The analysis code of this work is available on Github (https://github.com/MelissaSaichi/Tfh_GMCSF-DC).

## Acknowledgments

We would like to thank the NGS platform of Institute Curie for generating the single-cell RNA sequencing data, the Genomics Facility of Sanofi-Boston for generating the transcriptomic data and the technology platform of IRSL for their critical assistance in cell sorting and Flow cytometry analysis. We wish to thank also Dr. Jasna Medvedovic and Dr. Cristina Ghirelli for her critical review on the data and manuscript.

## Supplementary Information for

### Supplementary Materials and Methods

#### Human subjects

Apheresis blood from healthy human blood donors were obtained from Etablissement Francais du Sang (French Blood Establishment) after written informed consent and in conformity with Institute Curie and Research Institute Saint-Louis ethical guidelines. Gender identity and age from anonymous donors were not available, but all donors were between 18 and 70 years old (age limits for blood donation in France).

#### Blood Dendritic Cell Purification

Peripheral blood mononuclear cells (PBMC) were isolated by centrifugation on a density gradient (Lymphoprep, StemCell Technologies) following standard protocols. Primary blood DC were purified according to an established protocol (37). In brief, total PBMC were enriched in DC using the EasySep Human Pan-DC Pre-Enrichment kit (StemCell Technologies). Enriched DC were then sorted to obtain 98% purity on an MoFloAstrios Cell Sorting (Beckman Coulter) or FACS ARIA III (BD Technologies), as Lineage- (CD3, CD14, CD16, CD56, CD20 and CD19) (Miltenyi Biotech), CD4+ (Biolegend), CD11c+ (BioLegend), CD1c+ (eBioscience).

#### Naive CD4^+^ T Lymphocytes purification

After enrichment from total PBMC using the CD4^+^ T cell isolation kit (StemCell Technologies), naive CD4^+^ T cells were magnetically isolated. Purity was at least 95%.

#### Flow cytometry analysis

Antibodies were titrated on the relevant human PBMC population and matched isotypes controls were used at the same final concentrations. For intracellular cytokine staining, CD4^+^ T cells were stimulated with 100ng/ml PMA plus 500 ng/ml Ionomycin plus Brefeldin A (eBioscience) for 4 hours. When cells were sorted before intracellular staining, they were cultured overnight in X-VIVO medium at 10^6^ cells/ml before PMA and Ionomycin stimulation. To exclude dead cells, CD4^+^ T cells were stained using the LIVE/DEAD Fixable yellow dead cell stain kit, following manufacturer’s instructions (ThermoFischer Scientific). Cells were fixed and permeabilized using the IC Fix and Permeabilization buffers (eBioscience). Intracellular cytokines were revealed with fluorescently conjugated antibodies against IL-21 (Biolegend), TNF-□ (BioLegend), IL-4 (eBioscience), IFN-γ (eBioscience), and IL-17A (RD Technologies) or matched isotype controls (eBioscience) and acquired on a LSR Fortessa instrument (BD). For transcription factor intra-nuclear staining, dead cells were first stained with a yellow dye (BioLegend), followed by PD1 (Biolegend) and CXCR5 (BD) staining. After fixation and permeabilization using the FOXP3 IC buffer kit (eBioscience), cells were stained with an anti-BCL6 antibody (BD), TBET, GATA3, RORC, C-MAF, or SAP antibodies (eBioscience) and acquired on a LSR Fortessa instrument. As a control for intracellular staining of transcription factors, cells were stained with matched isotype controls at the same concentration as the transcription factor antibodies. The fluorescence obtained in each channel and in each population in the presence of the isotype control antibody (Fluorescence minus one [FMO]) was subtracted from the fluorescence obtained by the specific staining of transcription factors in each population.

#### T/B co-culture

After 4 days of co-culture with total GM-CSF-DC, CD4^+^ T cells were FACS sorted as PD-1^high^CXCR5^+^ (T^fh-like^ cells) or PD-1^low^CXCR5^+^ (T^low^ cells). Allogeneic PBMC were thawed and, after a round of human memory B cell Enrichment, memory B cells were magnetically sorted using the EasySep Human Memory B cell isolation kit (StemCell Technologies). T and B cells were co-cultured in X-VIVO medium in round-bottom plates (2.5 × 10^5^ T cells and 2.5 × 10^5^ memory B cells). At day 10 of culture, cells were harvested for FACS analysis.

#### Quality Control and pre-processing of expression matrices

We performed single cell RNA sequencing analysis using Chromium 10X technology of naive CD4^+^ T cells stimulated with Medium-DC, LPS-DC or GM-CSF-DC. Cell Ranger Software was used to generate fastq files and align them on Grch38 human reference genome. The expression matrix datasets were loaded on R (version 4.0.0) and the whole analysis was performed using Seurat package (version 3.2.0 https://github.com/satijalab/seurat). The three datasets corresponding to each condition were analyzed separately. For each sample, cells expressing at least 100 genes were kept to discard debris cells. Pre-processing steps were applied to remove cells with less than 100 expressed genes or having more than 20% of mitochondrial transcripts. Upper cutoffs of 8,000 and 90,000 were manually set for the nFeatures and nUMI respectively for each sample. In contrast, lower cutoffs of 3,500 and 20,000 were also set for the nFeatures and nUMI, respectively. Normalization to 10,000 reads, centering, and scaling were sequentially applied on the expression matrices to correct for the sequencing depth variability.

#### Downstream analysis

We used MAGIC (Markov Affinity-based Graph Imputation of Cells) (https://github.com/KrishnaswamyLab/MAGIC) imputation method to denoise the data and correct for the dropouts, and stored the corrected expression matrices into the “imputed” slot of the Seurat objects. To construct Th-related gene signatures, we used “AddModuleScore” Seurat function using 10 control genes and 4 bins. Cells, which overexpressed (or downexpressed) a given module, were attributed positive (or negative) scores. This strategy allowed us to point out cells, which co-express the genes, used to identify cell types.

The analysis code is deposited on Github: https://github.com/MelissaSaichi/Tfh_GMCSF-DC.

#### Dimension reduction and Clustering

The normalized count matrices were used to identify the highly variable genes within each dataset separately using the “mvp” method implemented in the “FindVariableFeatures” function. For each sample, PCA dimension reduction was applied on the top 3,000 genes, and the first 20 principal components (PC) were used for further steps, including cell community detection, clustering and non-linear dimension reduction. Cell clusters were identified using a shared nearest neighbor (SNN) clustering algorithm, which consists in a calculation of the k-nearest neighbors (k=30) then identification of cell communities (clusters) with a resolution parameter of 0.4. Non-linear dimensionality reduction methods, Uniform Manifold Approximation and Projection (UMAP) was used to explore and visualize the datasets, given as input the top 20 principal component genes.

#### Imaging flow cytometry

Samples were acquired with ImageStreamX MkII technology (Amnis/Luminex). Lasers power were set as 25mW for the 405 nm, 80 mW for the 488 nm and 200 mW for the 561 nm to excite respectively DAPI viability dye in channel 7 (430–505 nm filter), anti-CD40 FITC in channel 2 (480– 560 nm filter) and anti-ICOS-L-PE in channel 3 (560–595 nm filter). Channels 1 (430–480 nm filter) and 9 (570–595 nm filter) were both used to collect bright fields. Data were acquired with the 60× magnifications, a 7 µm core size and low flow rate. Analysis was performed with IDEAS software to calculate Brightfield circularity (arbitrary unit) for each live population of interest selected based on their Mean Fluorescence Intensities of CD40 and ICOS-L.

#### Cytokine quantification

Cytokines were quantified in the supernatants using ELISA for IL-21 (BioLegend) and CXCL13 (R&D Systems) or CBA flex set for IL-2, IL-3, IL-4, IL-5, IL-9, IL-10, IL-13, IL-17A, GM-CSF, TNF, and IFN-γ (BD), following the manufacturer’s protocol.

#### Statistical analysis

Statistical analysis was performed using the Prism software v7 (GraphPad). Paired Wilcoxon or t test were applied to compare two groups. Mann-Whitney test was used for non-paired analysis. Significance was retained for P < 0.05. The asterisks in the figures show the statistical significance as compared to the condition without any asterisks, *P < 0.05, **P < 0.01, ***P < 0.001.

#### Data, software and code Availability

Software used for flow cytometry data analysis was FlowJo software (TreeStar).

Software used for CBA analysis was FCAP Array v3.

Software used for statistical analysis was Prism software v5 (GraphPad).

Software used for statistical analysis and modeling was R version 4.0.0 The R packages used to perform this study are: Seurat Package 3.2.0

This study did not generate code.

---

**Fig S1.**
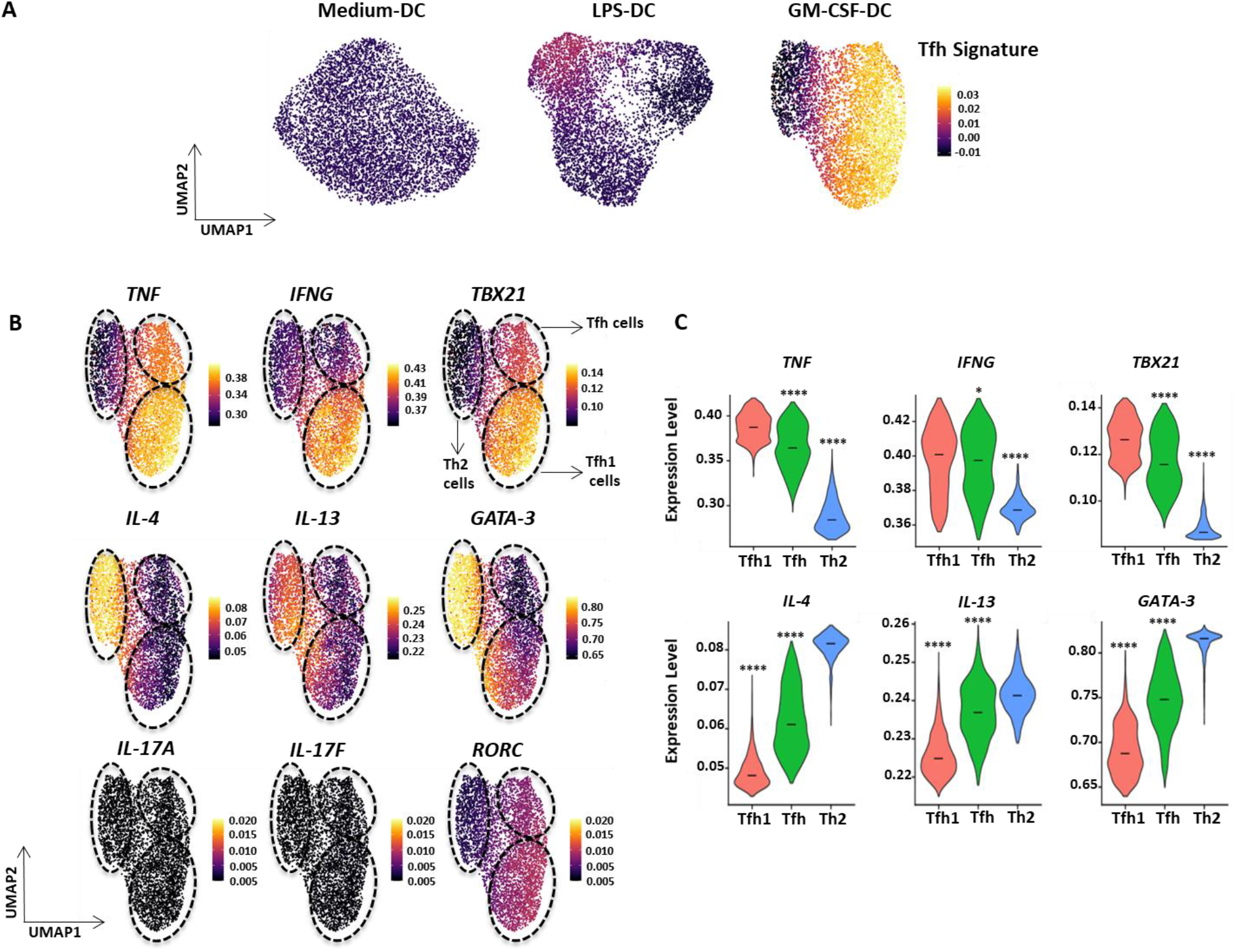
Expression of Th-related genes by CD4^+^ T cells activated by GM-CSF-DC, LPS-DC and Medium-DCType. **(A)** UMAP representation of the Tfh signature score of differentiated CD4^+^ T cells in the three following conditions of DC activation: Medium-DC, GM-CSF-DC and LPS-DC. Signature scores were calculated by the co-expression of the Tfh related genes (BCL6, PD-1, CXCR5, IL-21). **(B)** Corrected expression values of the Th1, Th2 and Th17 related genes respectively, represented on the UMAP plot of the CD4^+^ T cells differentiated by GM-CSF-DC. Th1 and Tfh related genes are co-expressed only by the CD4^+^ T cells differentiated by GM-CSF-DC condition. **(C)** Violin plot representation for the expression level of Th1 and Th2 related genes in Tfh (including pure-Tfh and Tfh1)- and Th2-enriched clusters. Asterisks above a violin plot show statistical differences as compared to the violin plot condition without any asterisks. *P < 0.05, **P < 0.01, ***P < 0.001

**Fig. S2.**
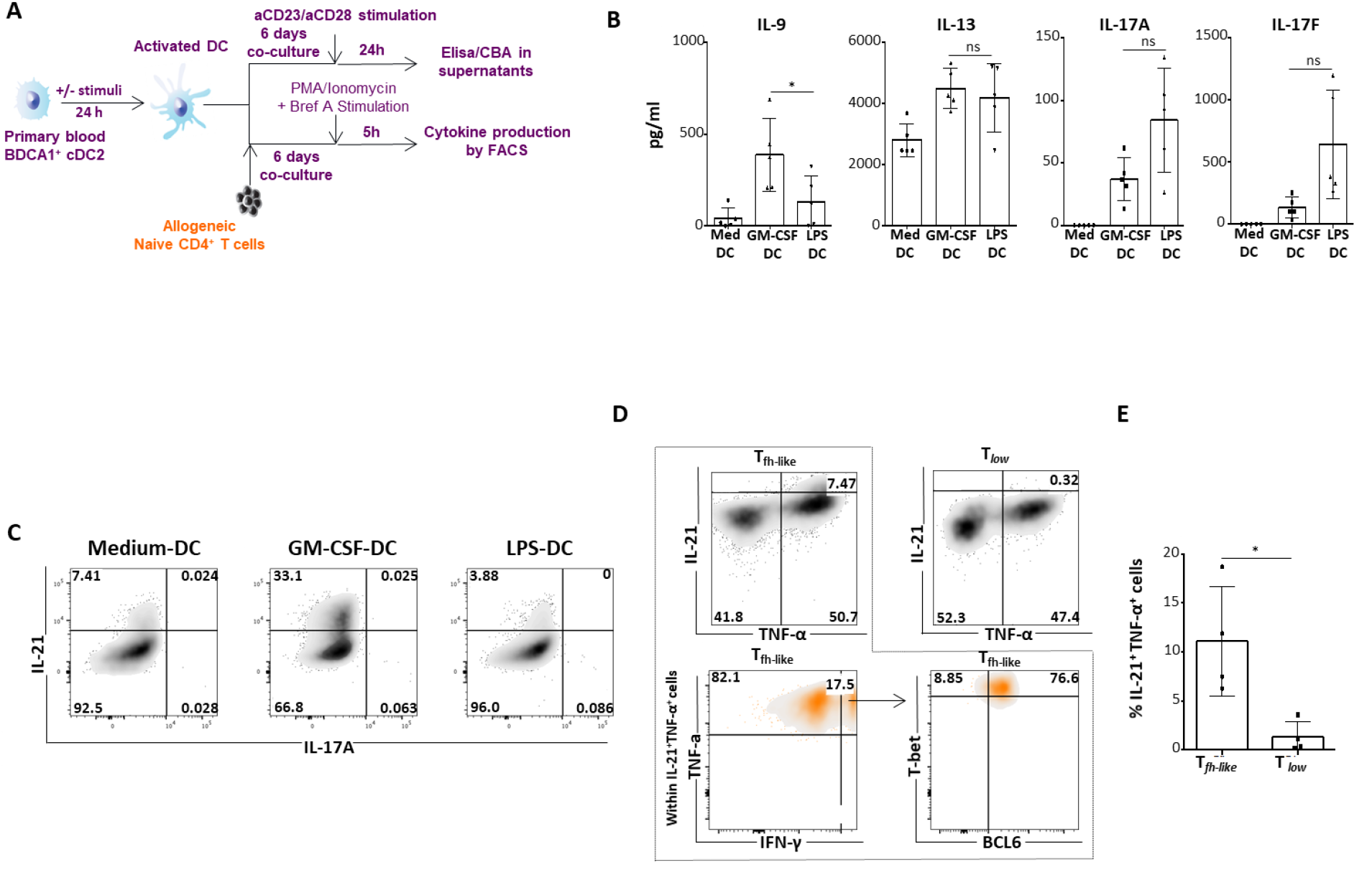
Cytokine profile of CD4^+^ T cells differentiated by GM-CSF-DC. **(A)** Experimental approach for the measurement of cytokines produced by CD4^+^ T cells differentiated by activated DC. DC were activated for 24h in the presence of GM-CSF, LPS or only Medium, followed by 6 days co-culture with allogeneic naive CD4^+^ T cells. At day 6, T cells were stimulated either with aCD3/aCD28 beads for additional 24h to measure cytokine secretion in the supernatants or with PMA/ionomycin and brefeldin-A for 4h, permeabilized and stained for the detection of several cytokines at a single cell level by flow cytometry. **(B)** CBA (IL-9, IL-13, IL-17A, IL-17F) assay for the measurement of cytokines in the supernatants of T cells in each condition of DC activation. Data are mean ± SEM from 5 independent experiments (n=5). **(C)** Intracellular staining for the expression of IL-21 and IL-17A by flow cytometry in T cells, one representative experiment (n=6). **(D)** Intracellular staining for the expression of IL-21, TFN-□, IFN-γ, BCL6 and T-BET by sorted T^fh-like^ and T^low^ cells in GM-CSF-DC condition for one representative experiment (n=4). **(E)** Percentages of IL-21^+^TNF-α^+^ cells from data as shown in D for four independent experiments (n=4), *, P < 0.05; **, P < 0.01; ***, P < 0.001, by Wilcoxon or Student’s t test.

**Fig. S3.**
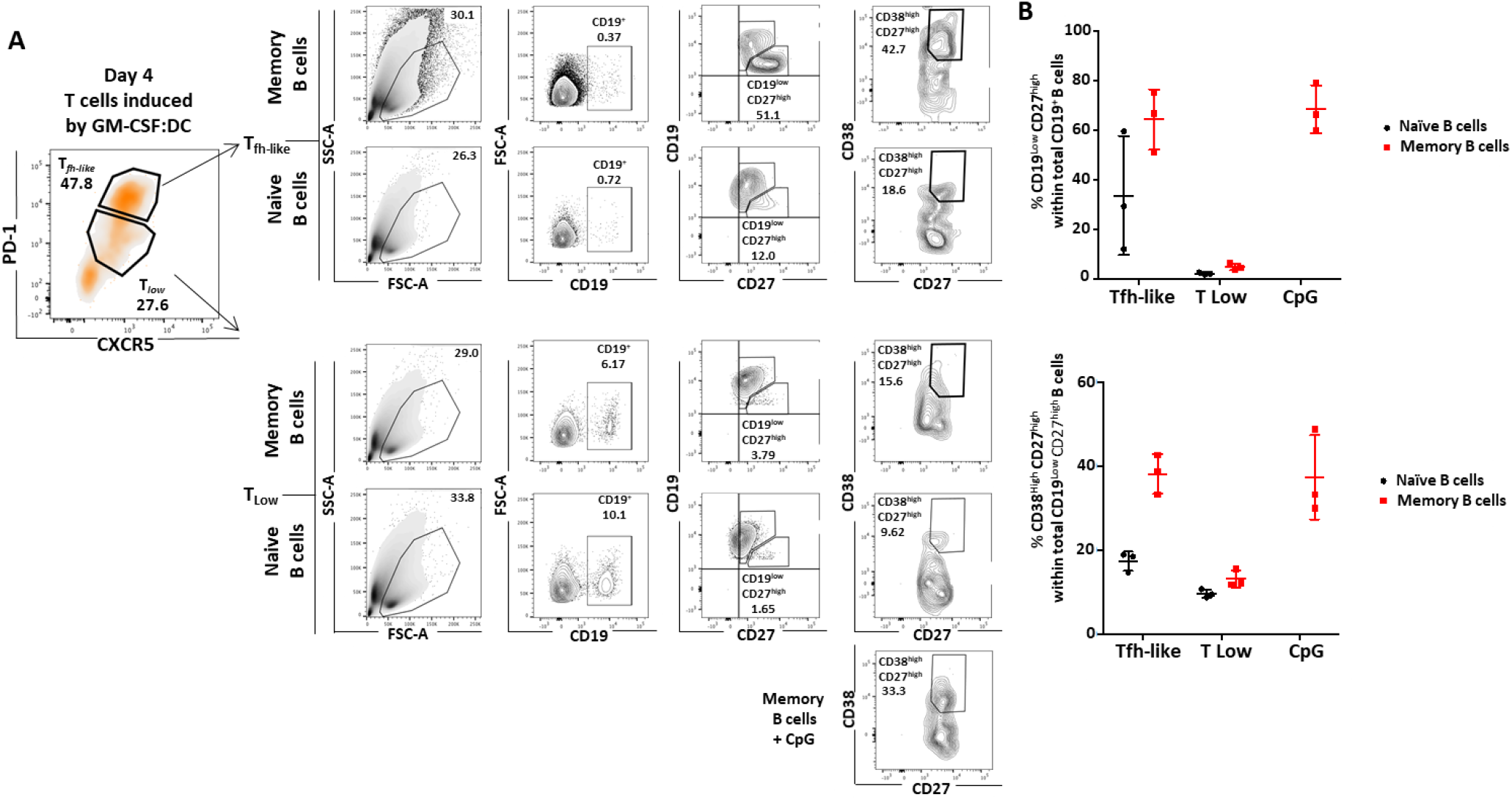
Plasma Cell differentiation induced by GM-CSF-DC-activated Tfh cells. (A) Tfh-like and T^low^ cells were sorted after four days of co-culture of naive CD4^+^ T cells and GM-CSF-DC based on the expression of PD-1 and CXCR5. T cells were then put in co-culture with allogeneic memory or naive B cells for 10 days. Extracellular staining for the expression of CD19, CD27 and CD38 for one representative experiment (n=3). **(B)** Percentages of CD19^low^CD27^high^ cells gated in total CD19^+^ cells and CD38^high^CD27^high^ gated in CD19^low^CD27^high^ cells from data as shown in A (n=3), *, P < 0.05; **, P < 0.01; ***, P < 0.001, by Wilcoxon or Student’s t test.

**Fig. S4.**
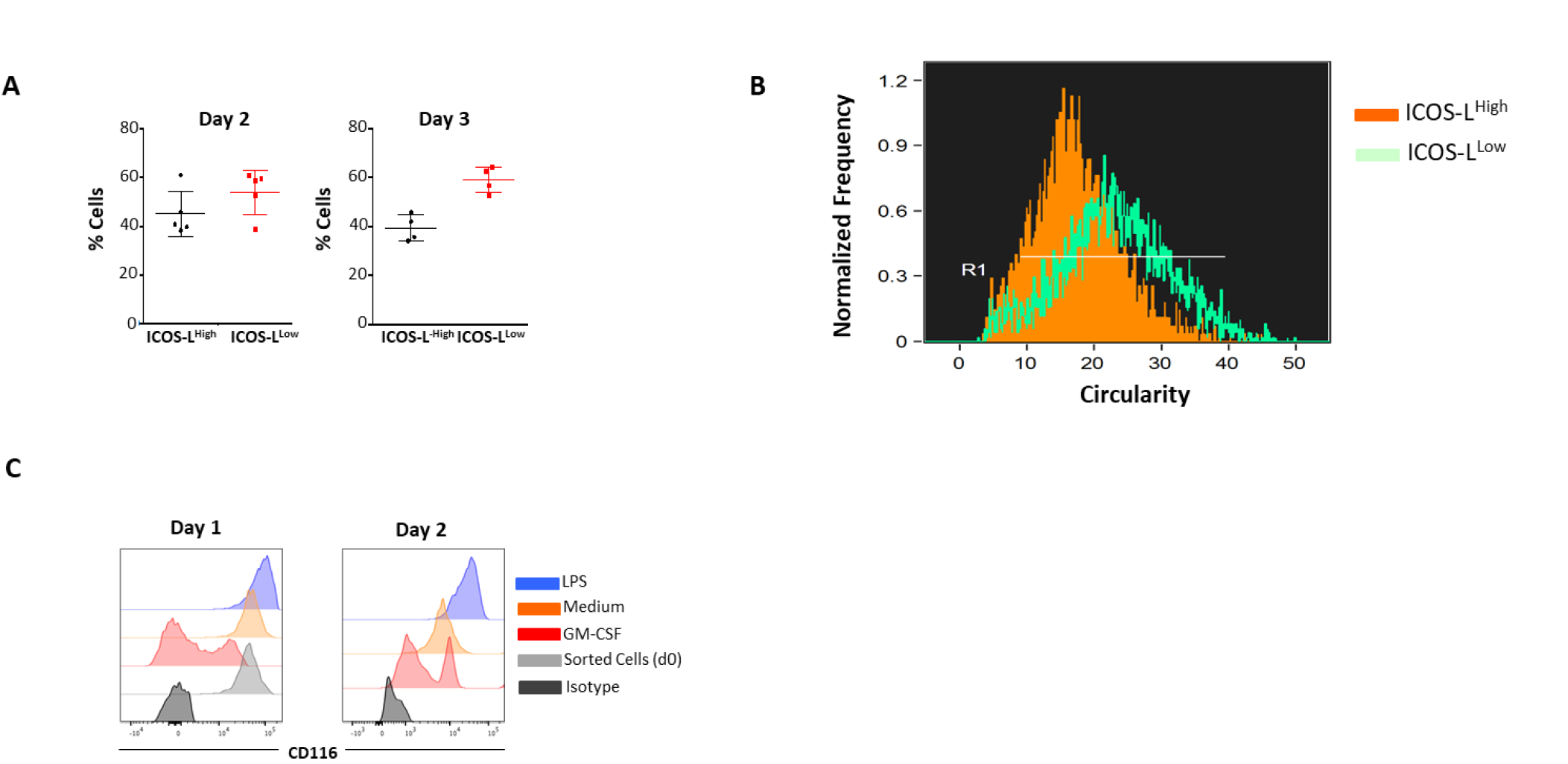
Morphological and phenotypic characterization of GM-CSF-DC. **(A)** Percentages of recovered ICOS-L^High^ and ICOS-L^Low^ DC after 2 and 3 days of activation with GM-CSF from data as shown in FIG 4D (n=5). **(B)** Histograms comparing the circularity levels of both ICOS-L^High^ and ICOS-L^Low^ GM-CSF-DC at day 2 by Imaging Flow Cytometry approach (n=3). **(C)** Expression of CD116 by FACS of human DC right after sorting (light grey histogram) or DC activated with LPS (blue histogram) or GM-CSF (red histogram) for 48h as compared to non-activated DC (orange histogram) and isotype control (dark grey histogram) for one representative experiment (n=4), *, P < 0.05; **, P < 0.01; ***, P < 0.001, by Wilcoxon or Student’s test.

**Fig. S5.**
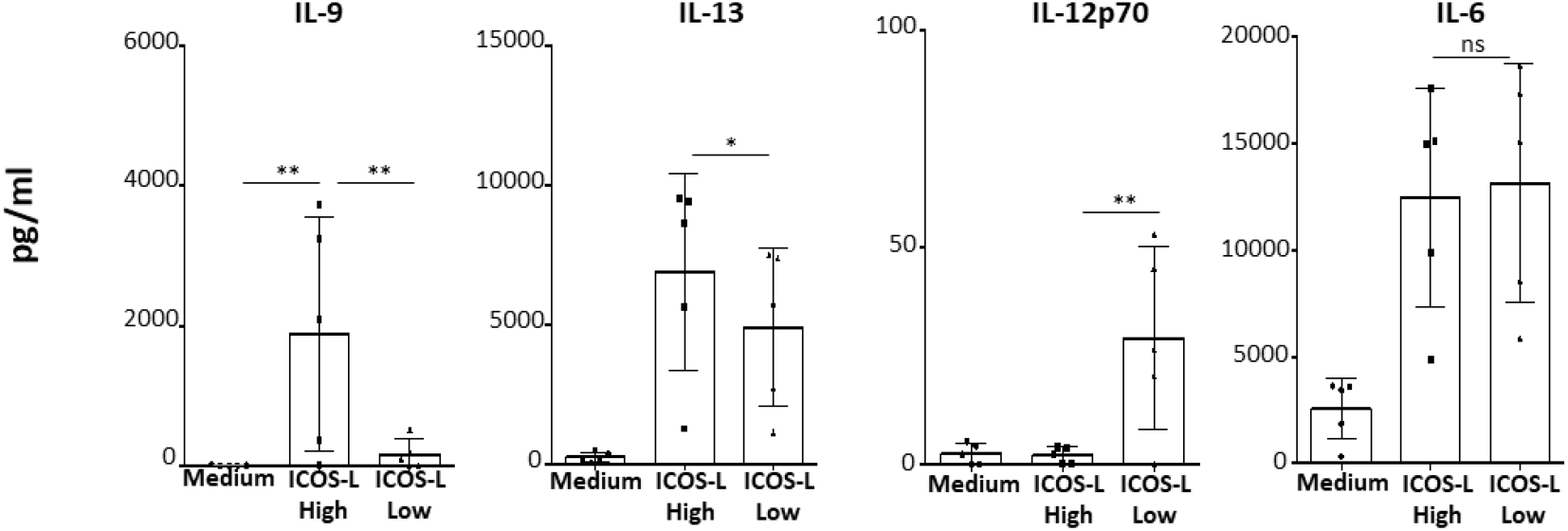
Cytokine profile of CD4^+^ T cells induced by either ICOS-L^High^ or ICOS-L^Low^ GM-CSF-DC. (A) CBA (IL-9, IL-13, IL-12p70, and IL-6) assay for the measurement of cytokines in the supernatants of CD4^+^ T cells differentiated either by ICOS-L^High^ or ICOS-L^low^ GM-CSF-activated DC, after additional 24h stimulation with aCD3/aCD28 beads. Data are mean ± SEM from 5 independent experiments (n=5), *, P < 0.05; **, P < 0.01; ***, P < 0.001, by Wilcoxon or Student’s test.

**Fig. S6.**
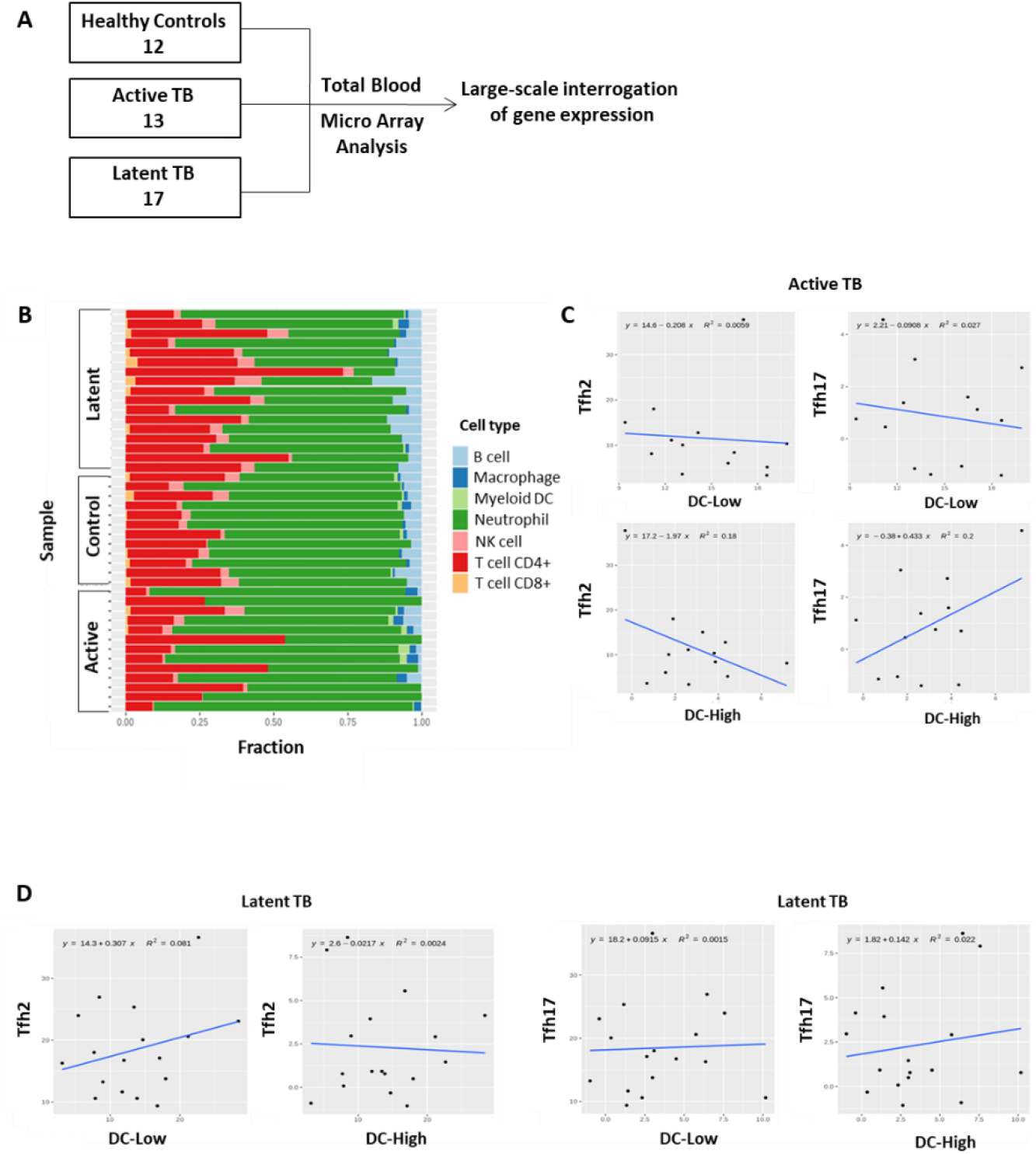
Correlation of in-house constructed DC-High and DC-low signatures with Tfh1 or Tfh2 cells in MTB patients. **(A)** Experimental design of the training set by Berry et al (GSE19904) to decipher the transcriptional signatures of tuberculosis patients. Microarray analysis was performed on Whole Blood (WB) samples collected from healthy controls (n=12), active tuberculosis patients (n=13) and latent TB patients (n=17). **(B)** Deconvolution of the Berry London training set revealed the presence of the major immune cell types. The deconvolution was performed using Quantiseq method from the immunedeconv R package. This method outputs the fractions of each cell type within each sample and therefore, allows inter sample and inter cell type comparisons. **(C)** and **(D)** Collinearity assessment of the corresponding signatures of Tfh2 and Tfh17 with the respective DC subtype signatures DC-Low and DC-High within active and latent TB patients respectively. Each dot represents a sample. A linear regression model was applied to fit the dots repartition.

**Fig. S7.**
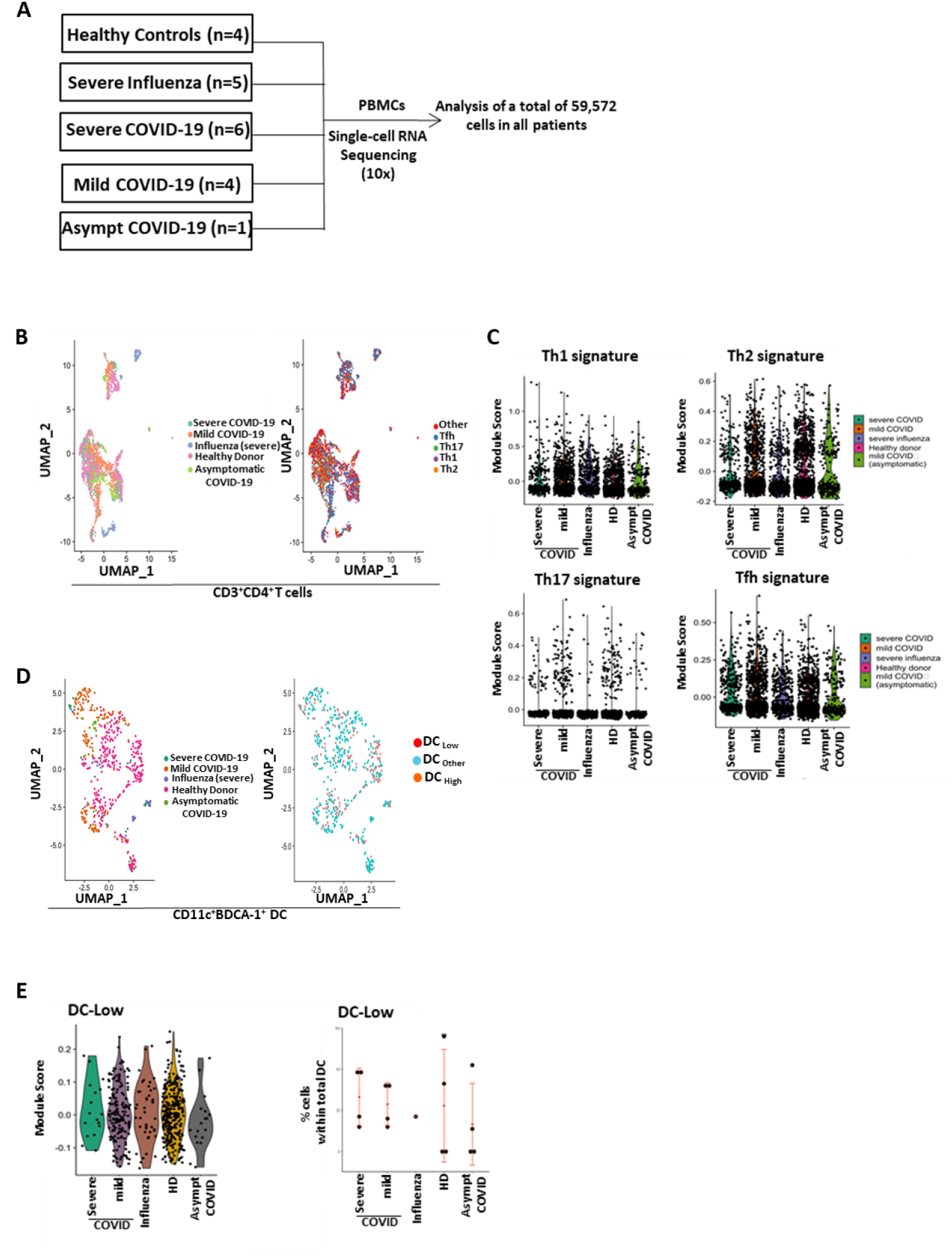
Tfh1 cells are positively correlated with the presence of DC-Low signature in mild COVID-19 patients. **(A)** Experimental design to decipher correlation between DC and Tfh signatures in COVID-19 patients. scRNAseq analysis was performed in PBMC from healthy controls or active COVID-19 (mild and severe) and severe influenza patients. **(B)** UMAP representation of CD3^+^CD4^+^ T cells in all patient groups (left) as well as of different T-helper signatures (right). **(C)** Violin plot representation for the expression level of Th1-, Th2-, Th17- and Tfh-related signatures in all patient groups. **(D)** UMAP representation of CD11c^+^BDCA1^+^ DC in all patient groups and representation for the distribution of cells expressing signatures for ICOS-L^Low^ (DC-Low) and ICOS-L^High^ (DC-High) GM-CSF-activated DC based on the top 10 genes detected by the transcriptomic analysis. **(E)** Violin plot representation for the expression level of DC-Low signature in all disease groups as well as the percentages of cells positive for the DC-Low signature among the total DC detected in each patient group.

**Table S1.** Molecular signatures used for the identification of Tfh, Th1, Th2 and Th17 cells.

| Tfh | Th1 | Th2 | Th17 |
| --- | --- | --- | --- |
| BCL6 | TNF | IL4 | IL17A |
| CXCR5 | IFNG | IL5 | IL17F |
| PDCD1 | TBX21 | GATA3 | RORC |
| IL21 |  |  |  |

Tfh: T follicular helper cells; BCL6: B-cell lymphoma 6 protein; CXCR5: C-X-C chemokine receptor type 5; PDCD1: Programmed cell death 1 (PD-1); IL21: Interleukin-21; Th1: T helper 1 cells; TNF: Tumor necrosis factor; IFNG: Interferon Gamma; TBX21: T-box transcription factor (T-BET); Th2: T helper 2 cells; IL-4: Interleukin-4; IL-5: Interleukin-5; GATA3: GATA binding protein 3; Th17: T helper 17 cells; IL17A: Interleukin-17A; IL17F: Interleukin-17F; RORC: RAR related orphan receptor C.

**Table S2.** Molecular signatures used for the identification of Tfh, Th1, Th2, Th17, Tfh1, Tfh2, and Tfh17 cells.

| Tfh | Th1 | Th2 | Th17 | Tfh1 | Tfh2 | Tfh17 |
| --- | --- | --- | --- | --- | --- | --- |
| ICOS | TNF IFNG | IL4 | IL17A | ICOS | IL4 | ICOS |
| PDCD1 | TBX21 | IL5 | IL17F | PDCD1 | IL5 | PDCD1 |
| CXCR5 |  | GATA3 | RORC | CXCR5 | GATA3 | CXCR5 |
| IL21 |  |  |  | IL21 | ICOS | IL21 |
| BCL6 |  |  |  | BCL6 | PDCD1 | BCL6 |
|  |  |  |  | TNF | CXCR5 | IL17A |
|  |  |  |  | TBX21 | IL21 | IL17F |
|  |  |  |  | IFNG | BCL6 | RORC |
|  |  |  |  | CXCR3 |  | CCR6 |

Tfh: T follicular helper cells; ICOS: Inducible T Cell costimulator; PDCD1: Programmed cell death 1 (PD-1); CXCR5: C-X-C chemokine receptor type 5; IL21: Interleukin-21; BCL6: B-cell lymphoma 6 protein; Th1: T helper 1 cells; TNF: Tumor necrosis factor; IFNG: Interferon Gamma; TBX21: T-box transcription factor (T-BET); Th2: T helper 2 cells; IL-4: Interleukin-4; IL-5: Interleukin-5; GATA3: GATA binding protein 3; Th17: T helper 17 cells; IL17A: Interleukin-17A; IL17F: Interleukin-17F; RORC: RAR related orphan receptor C; Tfh1: type 1 T follicular helper cells; CXCR3: C-X-C Motif Chemokine Receptor 3; Tfh2: type 2 T follicular helper cells; Tfh17: type 17 T follicular helper cells; CCR6: C-C Motif Chemokine Receptor 6.

**Table S3.**
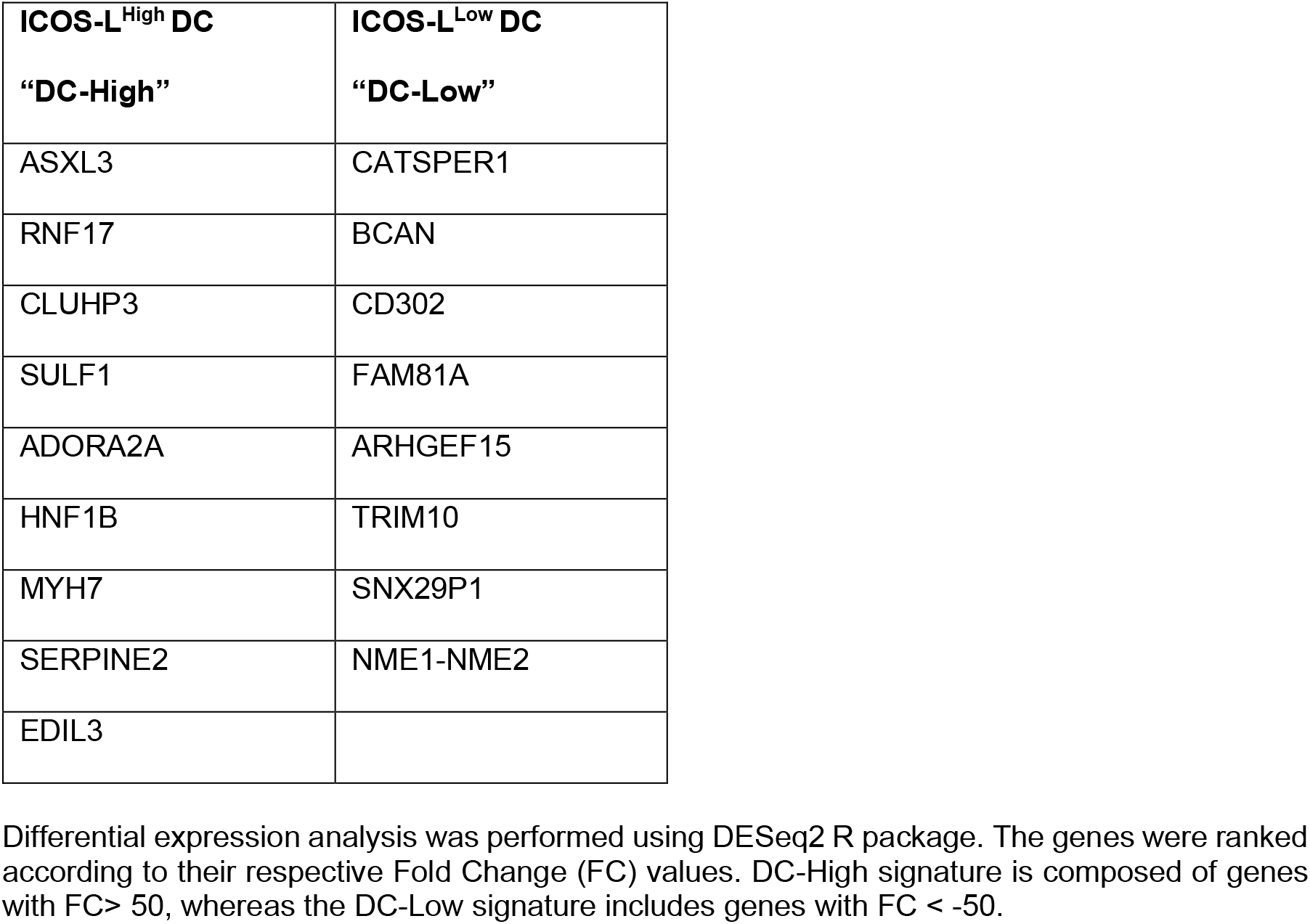
Top-9 differentially expressed genes characterizing the two in-vitro generated GM-CSF-activated DC subsets; ICOS-L^High^ (DC-High) and ICOS-L^Low^ (DC-Low).

| ICOS-L<sup>High</sup> DC<br>“DC-High” | ICOS-L<sup>Low</sup> DC<br>“DC-Low” |
| --- | --- |
| ASXL3 | CATSPER1 |
| RNF17 | BCAN |
| CLUHP3 | CD302 |
| SULF1 | FAM81A |
| ADORA2A | ARHGEF15 |
| HNF1B | TRIM10 |
| MYH7 | SNX29P1 |
| SERPINE2 | NME1-NME2 |
| EDIL3 |  |

Differential expression analysis was performed using DESeq2 R package. The genes were ranked according to their respective Fold Change (FC) values. DC-High signature is composed of genes with FC > 50, whereas the DC-Low signature includes genes with FC < -50.

